# Functional connectivity gradients as a common neural architecture for predictive processing in the human brain

**DOI:** 10.1101/2021.09.01.456844

**Authors:** Yuta Katsumi, Nada Kamona, Jiahe Zhang, Jamie G. Bunce, J. Benjamin Hutchinson, Mathew Yarossi, Eugene Tunik, Karen S. Quigley, Bradford C. Dickerson, Lisa Feldman Barrett

**Author notes:** **Correspondence** Yuta Katsumi, PhD, Department of Neurology, Massachusetts General Hospital, 149 13^th^ St, Charlestown MA 02129.

## Abstract

Predictive processing is emerging as a common computational hypothesis to account for diverse psychological functions subserved by a brain, providing a systems-level framework for characterizing structure-function relationships of its distinct substructures. Here, we contribute to this framework by examining gradients of functional connectivity as a low dimensional spatial representation of functional variation in the brain and demonstrating their computational implications for predictive processing. Specifically, we investigated functional connectivity gradients in the cerebral cortex, the cerebellum, and the hippocampus using resting-state functional MRI data collected from large samples of healthy young adults. We then evaluated the degree to which these structures share common principles of functional organization by assessing the correspondence of their gradients. We show that the organizing principles of these structures primarily follow two functional gradients consistent with the existing hierarchical accounts of predictive processing: A *model-error* gradient that describes the flow of prediction and prediction error signals, and a *model-precision* gradient that differentiates regions involved in the representation and attentional modulation of such signals in the cerebral cortex. Using these gradients, we also demonstrated triangulation of functional connectivity involving distinct subregions of the three structures, which allows characterization of distinct ways in which these structures functionally interact with each other, possibly subserving unique and complementary aspects of predictive processing. These findings support the viability of computational hypotheses about the functional relationships between the cerebral cortex, the cerebellum, and the hippocampus that may be instrumental for understanding the brain’s dynamics within its large-scale predictive architecture.

## Introduction

Predictive processing is emerging as a common neurocomputational hypothesis to account for diverse psychological functions subserved by a brain ^1,2^. A variety of specific proposals abound, but they are united by three hypothesized components that are thought to be implemented in a hierarchical arrangement within the brain’s architecture: (i) Prediction signals that the brain constructs using memory, also variously referred to as a generative or an internal model ^3^, “top-down” processing ^4–6^, a forward model ^7–9^, and “feedback” ^10^; (ii) incoming sense data from the body’s sensory surfaces encoded as the differences from predicted sensory inputs, called prediction errors, “bottom-up” processing, an inverse model, and “feedforward” signals; and (iii) precision signals that modulate predictions and prediction errors, corresponding to various attention signals ^11,12^. Prediction errors are potential teaching signals, but their capacity to update the model is thought to depend on how they are weighted by predicted precision signals, which are interpreted as the value of the information they provide, or “salience” ^13^ (see ^14^ for a discussion of precision and salience). Prediction signals are also thought to be weighted by their estimated value to explain the incoming sense data, similarly weighted by precision signals ^11–13^.

To date, predictive processing hypotheses have been offered to describe the computational capacities of several structures within the vertebrate brain, including the cerebral cortex ^1,6,13,15–19^, the cerebellum ^7–9,20^, and the hippocampus ^21–24^. Integrating these hypotheses into a systems-level framework for understanding brain structure-function relationships has the potential to computationally unify a variety of psychological and biological phenomena that are typically studied separately. In this paper, we contribute to this framework by examining gradients of functional connectivity, which reduce the dimensionality of complex brain connectivity data to describe continuous transitions in connectivity within a given structure and potentially offer parsimonious organizing principles to describe structure-function relationships ^25,26^. Intrinsic brain networks are often described in terms of patterns or modes of activity that reflect the similarity between the activity in different regions (i.e., functional connectivity). Functional connectivity gradients take this one step further by examining the patterns based on the similarity—not between activity—but between functional connectivity. Specifically, we investigated the degree of correspondence in functional connectivity gradients across the cerebral cortex, the cerebellum, and the hippocampus to test the hypothesis that these structures share common principles of functional organization that are consistent with the current hierarchical predictive processing accounts. Characterizing the coordination of functional connectivity gradients could be useful for formulating novel hypotheses about the role of these structures in the brain’s internal model of the body in the world, all in the ultimate service of predictive regulation of the body (i.e., allostasis) ^27^.

To identify such connectivity gradients, previous research capitalized on intrinsic functional connectivity derived from functional magnetic resonance imaging (fMRI) data collected when the brain is not being deliberately probed with an external task ^28,29^. Studies focusing on the cerebral cortex have most commonly revealed two gradients identifying gradual changes in connectivity profiles ^30–35^, which are consistent with the hypothesized role of different cortical areas in predictive processing ^36^. Previous research has also identified gradients that characterize the functional organization of the cerebellar cortex ^37,38^ and the hippocampus ^39–41^, suggesting that it may be possible to discover coordination in the connectivity gradients across these structures and the cerebral cortex. Functional coordination across different structures in the brain is also suggested by evidence describing learning systems across the cerebral cortex and cerebellum ^20,42^, cerebral cortex and hippocampus ^43,44^, and cerebellum and hippocampus ^45–49^. However, to our knowledge, no published study to date has examined how the functional organization of one structure relates to another, and what such functional correspondence might reveal about the contribution of these learning systems to the brain’s predictive processing architecture.

In the present study, we investigated the correspondence between intrinsic functional connectivity gradients in the cerebral cortex, the cerebellum, and the hippocampus using fMRI data collected at wakeful rest from large samples of healthy young adult participants. We derived functional connectivity gradients for each structure via diffusion map embedding, an established technique to nonlinearly reduce the dimensionality of large-scale connectivity data ^50,51^. We chose to examine our findings from the perspective of the cerebral cortex to consider how the cerebellar and the hippocampal gradients might align along the cerebral cortical gradients. We projected the cerebellar and hippocampal gradients onto the cerebral cortex to discover the common axes of functional organization. Finally, we performed a series of seed-based analyses of intrinsic functional connectivity to further characterize the functional correspondence between these gradients. This procedure allowed us to triangulate the three structures along a given connectivity gradient hypothesized to support particular aspects of predictive processing implemented in the brain. Finally, we evaluated the extent to which these findings were consistent with the computational hypothesis of predictive processing.

## Results

We examined the functional organization of the cerebral cortex, the cerebellum, and the hippocampus using fMRI data collected during wakeful rest from healthy young adult participants in the Human Connectome Project (HCP, *n* = 1,003) ^52^ as our primary sample and in the Brain Genomics Superstruct Project (GSP, *n* = 1,102) ^53,54^ as our validation sample. Following prior work on connectivity gradients ^32,38,41^, we analyzed the vertex-/voxel-wise similarity of intrinsic functional connectivity patterns in these structures based on the group average dense connectivity matrices via diffusion map embedding ^50,51^. Diffusion map embedding identifies multiple axes of variation in functional connectivity as continuous gradients, without ascribing each vertex or voxel to a singular functional unit (e.g., network). This feature allowed us to comprehensively examine the organizing principles of functional connectivity in the cerebral cortex, the cerebellum, and the hippocampus and evaluate their correspondence. Unless otherwise noted, all reported findings in the following sections were obtained using the HCP dataset. Overall similar results were obtained with the GSP dataset, with slight differences in the distribution of gradient values across the major subfields in the hippocampus (**Supplementary Fig. S2**). This is likely due to differences in the voxel resolution during data acquisition that had affected the ability to detect subfield specificity in the GSP dataset.

### Two principal functional gradients of the cerebral cortex consistent with a predictive processing account of brain function

To derive functional gradients of the cerebral cortex, we first constructed a subset of the whole-brain group average functional connectivity matrix with all cortical vertices, which we used as input to diffusion map embedding. From this analysis, we identified principal gradients that describe the maximal variance in functional connectivity patterns in the cerebral cortex, replicating those identified by prior work ^25,30,32,36^ (**Fig. 1** and **Supplementary Fig. S1**). These three gradients collectively explained >70% of variance in the data in both the HCP and GSP samples, with each accounting for >10%, as previously observed ^30^. Gradient 1 (G_1_) corresponded to a well-documented gradient consistent with cytoarchitectural evidence in the cerebral cortex ^55–57^. We refer to G_1_ as a *model-error* gradient, which we previously characterized as being anchored at one end by ensembles that can be described as initiating the prediction signals that constitute the brain’s internal model of its body in the world (e.g., default mode network), as well as those that estimate the precision of such signals (i.e., the model’s priors) (e.g., frontoparietal control network) ^13,36^. At the other end, this gradient was anchored by ensembles important for processing the sensory inputs that continually confirm or refine predictions, through supplying prediction errors (e.g., exteroceptive sensory networks) as well as those that estimate the precision of prediction error signals (e.g., salience network). Gradient 3 (G_3_) was consistent with a *model-precision* gradient that distinguishes between ensembles hypothesized to be involved in the representation of prediction and prediction error signals (e.g., default mode and exteroceptive sensory networks) and those involved in the implementation of attentional modulation—or precision—over those signals (e.g., salience, frontoparietal, and dorsal attention networks) ^13,36^. We also identified Gradient 2 (G_2_) replicating prior work, although its interpretation remains speculative. This gradient is anchored by ensembles primarily involved in the sensory representation of visual information at one end, and those involved in the representation of non-visual (somatosensory/motor, auditory, interoceptive) domains and multimodal integration at the other.

**Fig. 1.**
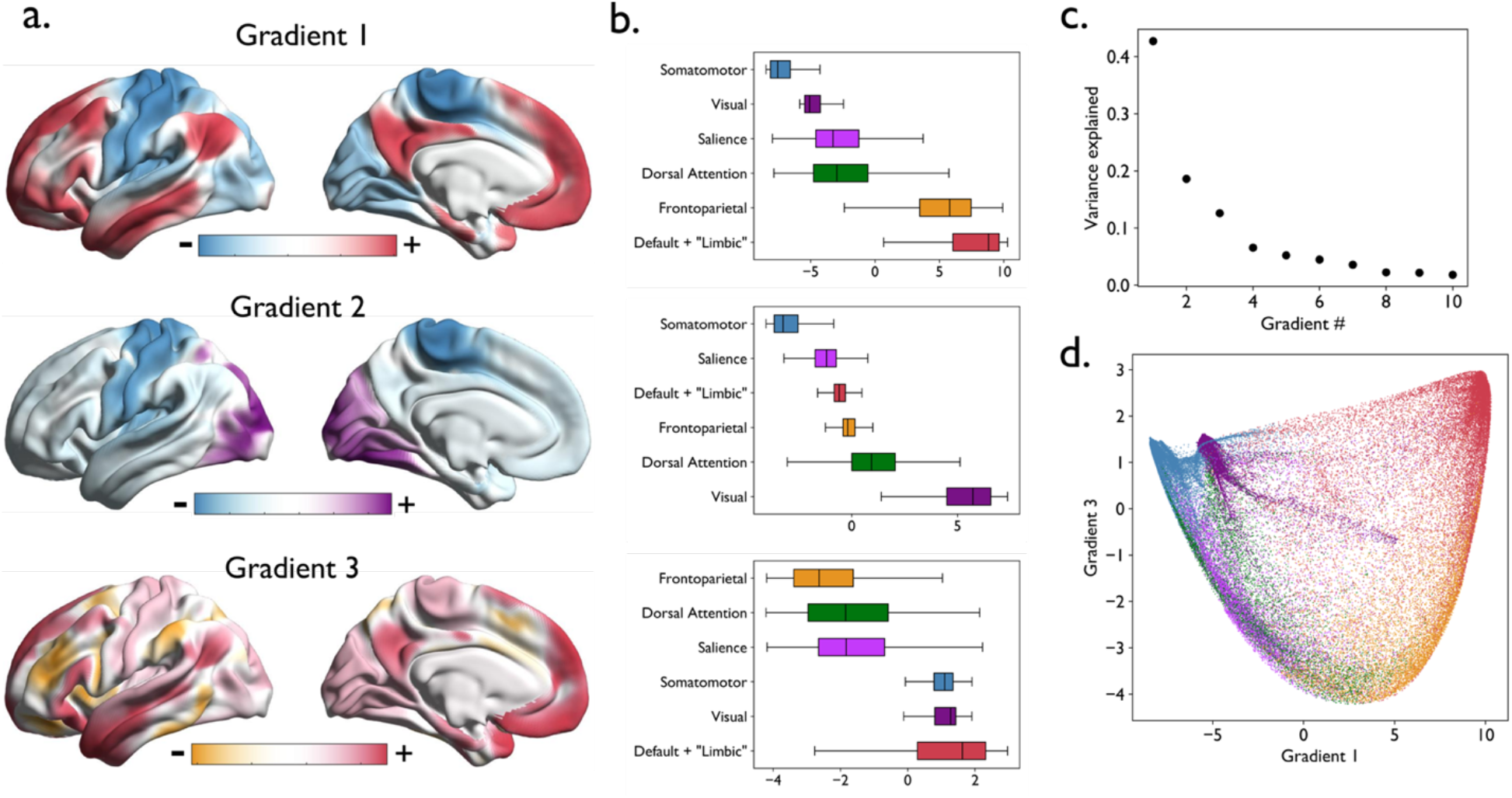
The principal gradients of human cerebral cortex based on the HCP data (*n* = 1,003). Functional connectivity gradients are a low dimensional spatial representation of connectivity profiles, such that the proximity of colors can be interpreted as greater similarity of connectivity patterns ^32^. (**a)** The three most dominant gradients of the cerebral cortex projected onto an inflated cortical surface (left hemisphere only). These gradients replicate previous findings and identify model-error (Gradient 1), visual-sensorimotor (Gradient 2), and model-precision (Gradient 3) gradients. **(b)** Box plots show the median and distribution of gradient values separately for each of the canonical functional network ^54^. The networks are ordered by the mean value. Vertices belonging with the so-called default mode and “limbic” networks are shown in the same color, as these networks are not always distinguished in the literature ^58^ and both contain agranular, limbic tissue ^17^. **(c)** A scree plot showing the proportion of variance explained by each of the ten gradients derived from diffusion map embedding. **(d)** A scatterplot depicting the relationship between Gradient 1 and Gradient 3, where each dot represents a cerebral cortical vertex color-coded by the corresponding network assignment.

### The principal functional gradients of the cerebellum and the hippocampus

To characterize the functional organization of the cerebellum and the hippocampus, we derived functional connectivity gradients separately for the cerebellum, left hippocampus, and right hippocampus. We first constructed group average functional connectivity matrices between all cerebellar voxels and all cortical vertices, as well as between all hippocampal voxels and all cortical vertices, which were used as input for diffusion map embedding. We derived cerebellar and hippocampal gradients by interrogating cerebello-cortical and hippocampo-cortical connectivity, respectively, rather than using connectivity within each structure, given our goal to characterize the functional organization of these structures in terms of their relation to the cerebral cortex. This approach is consistent with that of prior work examining functional gradients in the hippocampus and subcortical structures ^40,41,59–61^. The resulting gradients, therefore, represented the most dominant dimensions of spatial variability in functional connectivity patterns with the cerebral cortex within each structure.

In the cerebellum, we identified two gradients consistent with prior work on cerebellar gradients ^38^. Together, these gradients explained >60% of the variance in the data, with G_1_ accounting for >50% and G_2_ accounting for >10% in both the HCP and GSP samples. G_1_ captured a bilateral dissociation of lobules IV, V, and VI and lobule VIII from the posterior part of Crus I and II and the medial part of lobule IX, whereas G_2_ distinguished bilaterally the anterior parts of Crus I and Crus II along with lobule VIIb from the rest of the cerebellar cortex (**Fig. 2a** and **Supplementary Fig. S2a**). In the hippocampus, we also identified two gradients consistent with available evidence on hippocampal gradients ^41^. These gradients together explained >50% of the variance in the data within each hemisphere, with G_1_ accounting for >30% and G_2_ accounting for >20% of variance in both datasets. G_1_ generally captured spatial variation in functional connectivity along the longitudinal axis of the hippocampus, whereas the variation captured by G_2_ was observed in both the longitudinal axis and the transverse (i.e., medial-lateral) axis (**Fig. 2b** and **Supplementary Fig. S2b**). To properly understand G_2_ in terms of hippocampal microstructure ^41^, we performed Mann-Whitney tests to compare the distribution of gradient values for the major subfields within each hemisphere: Subicular complex, CA1-3, and CA4-dentate gyrus (CA4-DG), which were derived from the established segmentation protocol ^62^. This analysis revealed that G_2_ was anchored by the subiculum at one end and CA1-3 and CA4-DG at the other (*p* < .001), with no significant differences between the CA subfields. G_1_ better distinguished the CA subfields, with CA1-3 showing the highest values overall, followed by subiculum and then the CA4-DG (*p* ≤ .001).

**Fig. 2.**
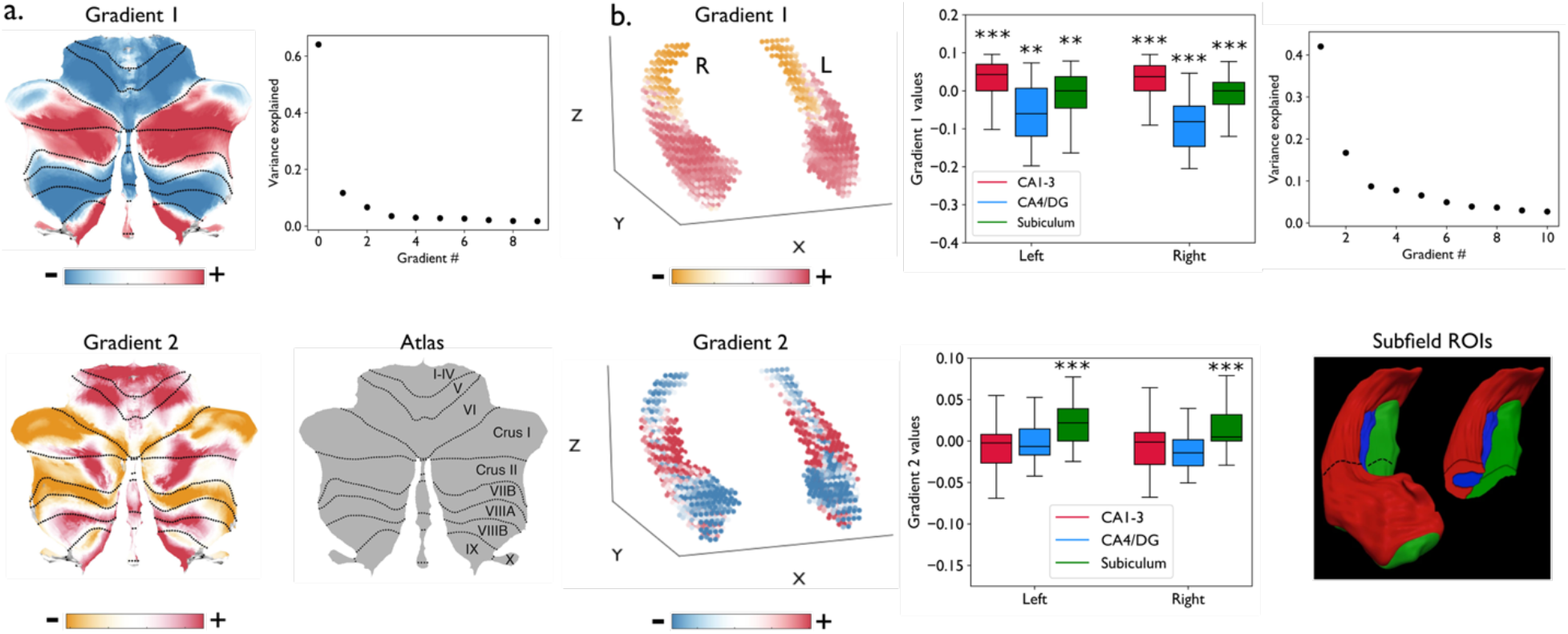
The principal gradients of human cerebellum and hippocampus based on the HCP data (*n* = 1,003). (**a)** The two most dominant gradients of the cerebellum replicated previous findings ^38^. **(b)** The most dominant gradient of the hippocampus replicated previous findings ^41^ and identified an anterior-posterior dissociation along the longitudinal axis, which also differentiated the major hippocampal subfields. The second most dominant gradient was also consistent with differences by hippocampal microstructure, with the subiculum exhibiting highest gradient values overall compared with the other two subregions. Box plots show the median and distribution of G_2_ values per subfields separately for each hemisphere. Asterisks denote significant (\*\*\**p* < .001, \*\**p* ≤ .001) differences relative to the other two subfields. DG = dentate gyrus. A figure illustrating the hippocampal subfields within the right hippocampus (red = CA1-3, blue = CA4-DG, green = subiculum) was reproduced from ^62^ with permission.

### Gradient-informed triangulation of intrinsic functional connectivity between the cerebral cortex, the cerebellum, and the hippocampus

Next, we calculated functional connectivity maps for the cerebral cortex weighted as a factor of gradient values for cerebellar and hippocampal gradients ^36,59^. This procedure allowed us to characterize these gradients in terms of their relations to the cerebral cortex. To calculate voxel-wise gradient-weighted functional connectivity with the cerebral cortex, we multiplied the group average cerebello-cortical (or hippocampo-cortical) functional connectivity matrix by a vector of voxel-wise values from cerebellar (or hippocampal) gradients. The weighted connectivity matrix was then reduced to a single cortical surface map representing the degree of connectivity between the cerebellum (or hippocampus) and the cerebral cortex along a particular gradient. For example, to characterize how cerebellar G_1_ relates to the cerebral cortex, we multiplied each row of the cerebello-cortical functional connectivity matrix (rows = cerebellar voxels, columns = cortical vertices) by the corresponding G_1_ value for that particular cerebellar voxel obtained from diffusion map embedding. In this way, the pattern of functional connectivity between each cerebellar voxel and all cortical vertices was weighted by its position on cerebellar G_1_. The G_1_-weighted cerebello-cortical connectivity values in this matrix were summed over all rows (i.e., all cerebellar voxels), resulting in a single cortical representation (projection) of this cerebellar gradient. We repeated this procedure for each gradient derived for the cerebellum and the hippocampus.

We then quantitatively assessed the correspondence between the cerebral cortical gradients (**Fig. 1** and **Supplementary Fig. S1**) and the gradient-weighted functional connectivity maps of the cerebellum (**Fig. 2a** and **Supplementary Fig. S2a**) and the hippocampus (**Fig. 2b** and **Supplementary Fig. S2b**) by computing vertex-wise Spearman’s rank correlations, while statistically controlling for autocorrelations ^63^. The model-error gradient in the cerebral cortex (G_1_) showed the strongest statistical correspondence to cerebellar G_1_ and hippocampal G_2_; the weighted functional connectivity maps of cerebellar G_1_ and hippocampal G_2_ showed moderate, statistically significant correspondence. The model-precision cortical gradient (G_3_) similarly showed strong correspondence to cerebellar G_2_ and hippocampal G_1_, with the weighted connectivity maps of cerebellar G_2_ and hippocampal G_1_ also showing strong correspondence (all *p*’s < .001) (**Fig. 3** and **Supplementary Fig. S3**).

**Fig. 3.**
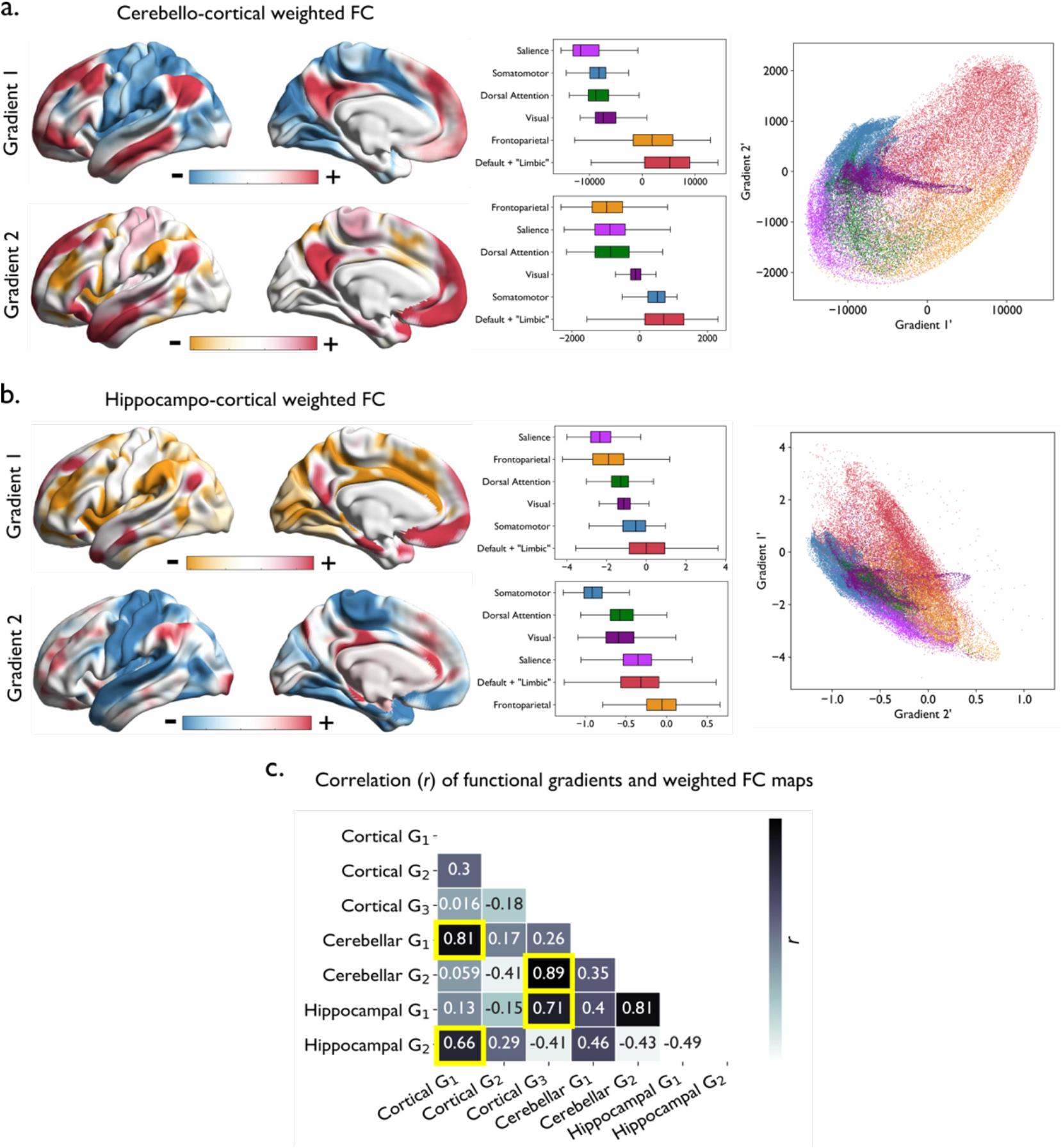
Gradient-weighted functional connectivity maps of the cerebellum and the hippocampus. **(a)** Cerebellar G_1_ captured a dissociation in functional connectivity most consistent with cortical G_1_, from ensembles including the default mode and frontoparietal networks to those including the exteroceptive sensory and salience networks (i.e., the model-error gradient). In contrast, cerebellar G_2_ captured a dissociation in connectivity most consistent with cortical G_3_, from ensembles including the default mode and exteroceptive networks to those including the frontoparietal and salience networks (i.e., the model-precision gradient). Box plots and scatterplots represent similar information as described in Fig. 1b, but here they are based on gradient-weighted functional connectivity maps. **(b)** Hippocampal G_1_ captured a dissociation in functional connectivity most consistent with cortical G_3_, whereas hippocampal G_2_ captured a dissociation in connectivity most consistent with cortical G_1_. **(c)** A similarity matrix illustrating the magnitude (Spearman’s *r*) of correlation between cortical gradients and gradient-weighted functional connectivity maps of the cerebellum and the hippocampus. Highlighted in yellow are the strongest cortico-cerebellar or cortico-hippocampal associations identifying the correspondence of gradients between a pair of structures (all *p*’s < .001). *p*-values associated with the entire correlation matrix is included as part of **Supplementary Fig. S5**). FC, functional connectivity.

A series of seed-based analyses of intrinsic functional connectivity between the three structures allowed us to further characterize the connectivity profiles for the cortical, cerebellar, and hippocampal gradients. This analysis was conducted separately for the cortical model-error gradient (G_1_) and its corresponding hippocampal (G_2_) and cerebellar (G_1_) gradients, and for the cortical model-precision (G_3_) and its corresponding hippocampal (G_1_) and cerebellar (G_2_), gradients (**Fig. 3c**). Seed regions of interest (ROIs) were defined as the vertices/voxels with the top and bottom 10% values on each gradient, resulting in two ROIs per structure per gradient. We then computed the mean BOLD activity time course based on all vertices/voxels within each ROI and correlated it with the time course of every vertex/voxel in the other two structures. If the connectivity gradients indeed correspond to one another across the three structures, then we should expect that the vertices/voxels representing the top (bottom) 10% of each gradient show stronger functional connectivity with each other than with other parts of these structures.

Results of this analysis based on the HCP data are summarized in **Fig. 4** (for results based on the GSP data, see **Supplementary Fig. S4**). **Fig. 4A-C** illustrate three of the seed ROIs used in this analysis, generated by identifying the vertices/voxels in each structure showing the top 10% of the gradient values along the cortical model-precision gradient as well as its corresponding cerebellar and hippocampal gradients (i.e., areas of each structure potentially more related to the representation of prediction and error signals). The same procedure was repeated with the vertices/voxels anchoring the bottom 10% of these gradients, yielding a set of ROIs representing the areas of each structure potentially more related to precision signals modulating prediction and error signals (**Fig. 4D-F**). In the hippocampus, the voxels anchoring the top and bottom 10% were exclusively localized to CA1-3 along the longitudinal axis. We defined the ROIs following the same procedure for the cortical model-error gradient and the corresponding cerebellar and hippocampal gradients, yielding a set of ROIs anchoring the top 10% of the vertices/voxels representing the areas of each structure potentially more related to the internal model (prediction signals) (**Fig. 4G-I**) as well as those ROIs anchoring the bottom 10% of the vertices/voxels representing the areas of each structure potentially more related to processing of sense data from the periphery (prediction error signals) (**Fig. 4J-L**). In the hippocampus, the voxels anchoring the top 10% (middle subregions) were localized to the subiculum medially and to CA1-3 laterally, whereas those anchoring the bottom 10% (anteroventral subregions) were predominantly localized to CA1-3.

**Fig. 4.**
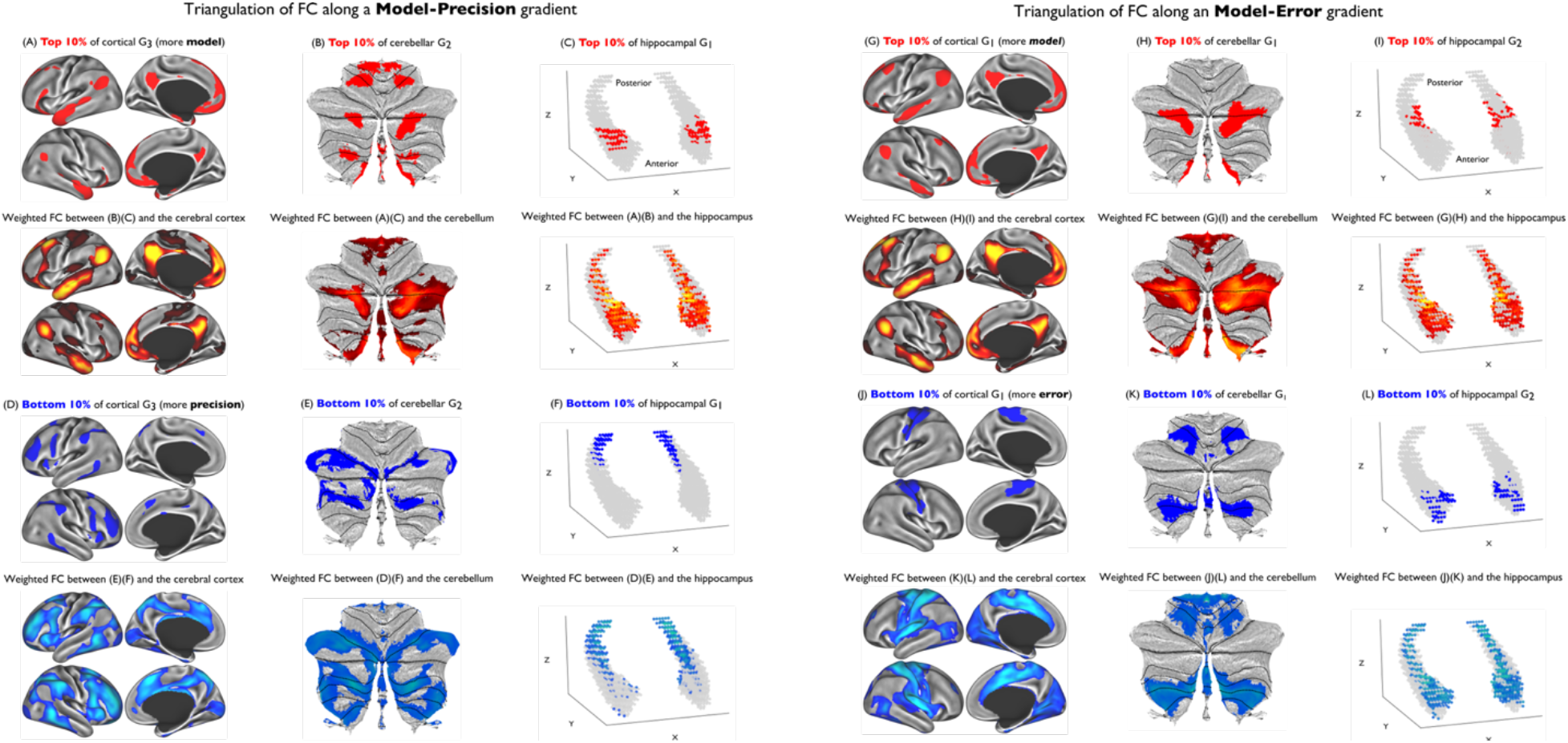
Triangulation of intrinsic functional connectivity between the cerebral cortex, the cerebellum, and the hippocampus along the cortical model-precision (left) and the cortical model-error (right) gradients in the HCP (*n* = 1,003) data. (A) Seed region of interest (ROI) representing the top 10% of the vertices in cortical model-precision G_3_. (B) Seed ROI representing the top 10% of the voxels in cerebellar G_2_. (C) Seed ROI representing the top 10% of the voxels in hippocampal G_1_. (D) Seed ROI representing the bottom 10% of the vertices in cortical model-precision G_3_. (E) Seed ROI representing the bottom 10% of the voxels in cerebellar G_2_. (F) Seed ROI representing the bottom 10% of the voxels in hippocampal G_1_. (G) Seed ROI representing the top 10% of the vertices in cortical model-error G_1_. (H) Seed ROI representing the top 10% of the voxels in cerebellar G_1_. (I) Seed ROI representing the top 10% of the voxels in hippocampal G_2_. (J) Seed ROI representing the bottom 10% of the vertices in cortical model-error G_1_. (K) Seed ROI representing the bottom 10% of the voxels in cerebellar G_1_. (L) Seed ROI representing the bottom 10% of the voxels in hippocampal G_2_. Functional connectivity maps shown here for a given structure were calculated through a combination of binarization and inclusive masking of the contributing maps as well as proportional thresholding (see **Methods**). We expect, and indeed observe, that there is remarkable spatial overlap between a given seed ROI (e.g., the top 10% of the vertices in cortical model-precision G_3_) and areas of the same structure functionally connected to parts of the other two structures anchoring the same end of the gradient (e.g., the top 10% of the voxels in cerebellar G_2_ and hippocampal G_1_).

The vertices representing the top 10% of the cortical model-precision gradient G_3_ (primarily those within the default mode network; **Fig. 4A**) showed relatively stronger positive functional connectivity with areas of the cerebellum overlapping with the top 10% of the voxels in cerebellar G_2_ (parts of lobule I-VI, posterior Crus I/II, and posterior lobule VIII/IX; **Fig. 4B**) and with areas of the hippocampus overlapping with the top 10% of the voxels in hippocampal G_1_ (dorsal anterior subregions; **Fig. 4C**), compared with other parts of the target structures. We identified similar patterns of spatial overlap in functional connectivity when using the top 10% voxels in cerebellar G_2_ and in hippocampal G_2_ as seed ROIs. Similarly, the vertices representing the bottom 10% of the cortical model-precision gradient G_3_ (primarily those within the salience and frontoparietal networks; **Fig. 4D**) showed relatively stronger positive functional connectivity with areas of the cerebellum overlapping with the bottom 10% of the voxels in cerebellar G_2_ (anterior parts of Crus I/II and lobule VIIb; **Fig. 4E**) and with areas of the hippocampus overlapping with the bottom 10% of the voxels in hippocampal G_1_ (posterior-most subregions), compared with other parts of the target structures.

We identified such pattern of functional connectivity triangulation similarly for the cortical model-error gradient and the corresponding cerebellar and hippocampal gradients, although there was overall less specificity in the connectivity pattern (**Fig. 4, right**). Specifically, the vertices representing the top 10% of cortical model-error G_1_ (primarily the default mode network; **Fig. 4G**) showed relatively stronger and positive functional connectivity with areas of the cerebellum overlapping with the top 10% of the voxels in cerebellar G_1_ (parts of Crus I/II and posterior lobule IX; **Fig. 4H**) and the areas of the hippocampus overlapping with the top 10% of the voxels in hippocampal G_2_ (middle lateral and medial subregions; **Fig. 4I**). Similarly, the vertices representing the bottom 10% of cortical model-error G_1_ (primarily the somatomotor network; **Fig. 4J**) showed relatively stronger positive functional connectivity with areas of the cerebellum overlapping with the bottom 10% of the voxels in cerebellar G_1_ (lobules IV, V, VI, and VIII; **Fig. 4K**) and the areas of the hippocampus overlapping with the bottom 10% of hippocampal G_2_ (ventral anterior subregions; **Fig. 4L**). Triangulation of functional connectivity appeared less specific along the model-error gradient, as the cortical and cerebellar subregions relevant for this gradient showed widespread connectivity throughout the hippocampus. These results point to the possibility that the distinct subregions of the cerebral cortex, the cerebellum, and the hippocampus form functional circuits that may contribute to the brain’s large-scale implementation of predictive processing. Importantly, we replicated this pattern of connectivity in a large, independent sample of healthy young adults (**Supplementary Fig. S4**), suggesting that our results are robust to variations in data acquisition parameters and preprocessing methods.

## Discussion

Accumulating evidence reveals that the organization of the cerebral cortex ^30,32–36,64–66^, the cerebellum ^38,59^, and the hippocampus ^39–41,67,68^ can be described with multiple gradients of structural and functional features in humans. In the cerebral cortex, converging evidence from network-, circuit-, and cytoarchitectural-levels of analysis suggest that such gradients can be interpreted as the organizing principles underlying predictive processing ^36,55,56,69,70^, guiding the flow of prediction signals, prediction error signals, and precision signals. In the present study, analyses of two large datasets, consisting of more than 2,000 participants, revealed that there are corresponding connectivity gradients across the cerebral cortex, the cerebellum, and the hippocampus, suggesting that these gradients might be meaningfully interpreted within a common computational framework. These results, and the specific computational hypotheses that they suggest, represent an important step toward an integrative account of brain function, building upon the existing literature on brain functional gradients that has so far largely focused on single regions without interrogating their interactions ^38,41,61,71^.

### Functional connectivity gradients as a common neural architecture for predictive processing

The principal cortical gradient was anchored, at one end, by ensembles that can be described as initiating the prediction signals that constitute the brain’s internal model of its body in the world (e.g., default mode network), as well as the ensembles that estimate the precision of such signals (e.g., frontoparietal control network). At the other end, this gradient was anchored by ensembles important for processing sensory inputs that continually confirm or refine the internal model (e.g., exteroceptive sensory networks) as well as those that estimate the precision of prediction error signals (e.g., salience network) (see ^13,36^ and references therein regarding the roles of cortical ensembles). Although in the present work we refer to this gradient as a model-error gradient, it has also been variably called by previous studies an internal-external gradient ^36^ and a transmodal-primary sensorimotor gradient ^32^. This gradient is consistent with a key structural hypothesis supported by more than 30 years of tract-tracing of distinctive cortico-cortical connections in mammalian brains ^55,57,72^ that describes the flow of prediction and prediction error signals throughout the cerebral cortex on the basis of cytoarchitectural properties (**Fig. 5**).

**Fig. 5.**
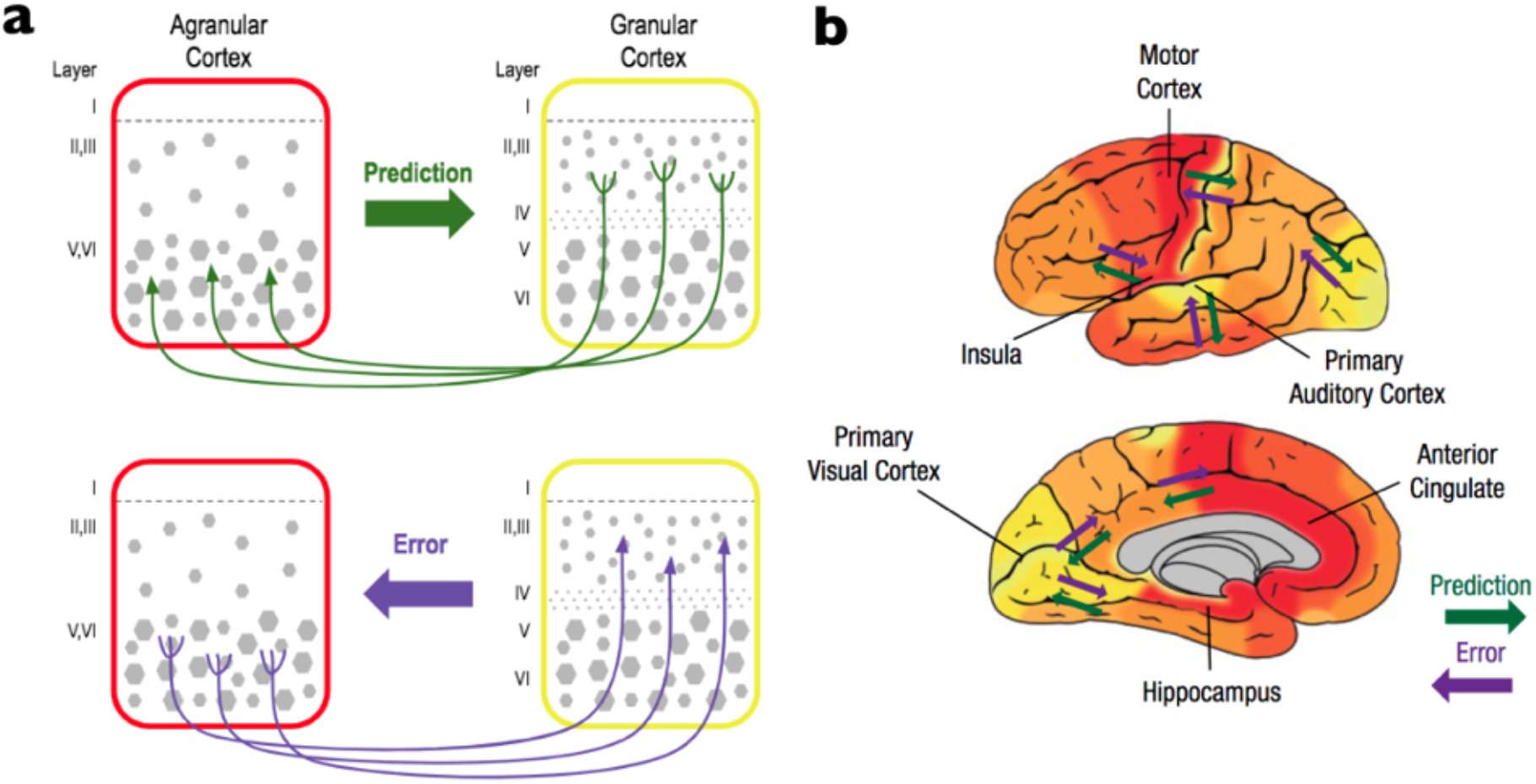
A gradient of predictive processing in the cerebral cortex. (a) Prediction signals originate in the deep layers (Layers V and VI) of less differentiated cortical areas (e.g., agranular cortex with undifferentiated Layers II and III and without a Layer IV) and terminate in superficial layers of areas with a more developed laminar structure (e.g., dysgranular cortices with differentiated Layers II and III and a rudimentary Layer IV or granular cortices with differentiated Layers II and III and a well-defined Layer IV). Prediction signals, as they cascade from agranular cortices to highly granular primary sensory areas, can be described as perceptual inferences that arise from the lossy compression that occurs during learning ^2,16^. Prediction error signals flow in the opposite direction, originating in the superficial layers (II and III) with more laminar differentiation and terminating in the deep layers (V and VI) of areas with less differentiated laminar architecture (66 as discussed in 2; see also recent work by 67, 68). This cytoarchitectural gradient is thought to support information compression in the cerebral cortex. Prediction errors are compressed and reduced in dimensionality ^13,73^ as they flow from the upper layers of highly granular primary sensory regions (whose upper layers contain many smaller pyramidal neurons with fewer connections) to less granular motor cortex ^74^ and other heteromodal regions (whose upper layers contain fewer but larger pyramidal neurons with many more connections), and finally to dysgranular and agranular limbic regions. (b) This structural model of cortico-cortical connections successfully predicts the flow of information in frontal, temporal, and parietal cortices in experiments with monkeys and cats ^55,56^. Figure adopted from ^2^, with permission.

Additional tract-tracing and optogenetic evidence support the involvement of this gradient in predictive processing. For instance, it has been shown that long-range connections exist between cortical limbic areas (e.g., anterior cingulate cortex) and primary sensory areas (e.g., V1) ^75^, which are two areas that anchor the ends of the model-error gradient, with the former thought to send sensory prediction signals to the latter ^76^. Such evidence is in line with other findings that a substantial fraction of activity in the visual cortex does not derive from incoming visual input per se ^77–81^, consistent with observations that the majority of synapses in V1 originate from top-down sources ^82^. Multimodal evidence also demonstrates the correspondence between the model-error gradient with regional variability in intracortical myelin ^83^ as well as cellular density and soma size ^33^, further substantiating the role of this gradient as a primary organizing principle in the cerebral cortex.

Recent research also describes this model-error gradient as important for learning and meaning-making. Unanticipated sense data (i.e., prediction errors), as they propagate from primary sensory regions (the external pole of this gradient) to agranular limbic regions (the internal pole), undergo lossy compression and are reduced in dimensionality ^13,16,73^. This process of information compression has also been described as conceptual learning ^13^ or construction of “generic priors” consisting of low-dimensional representations of the most frequent behavioral states ^84^. Such dimensionality reduction allows the brain to represent a large amount of information with a smaller population of neurons by decreasing redundancy and increasing efficiency, because smaller populations of neurons are summarizing statistical regularities in the spiking patterns of larger populations of neurons in the sensorimotor regions. Moving in the direction from internal to external poles, it is hypothesized that the low-dimensional, multimodal summaries are generatively reassembled as prediction signals in agranular limbic cortices, becoming successively decompressed and particularized as perceptual inferences (also called embodied simulations) that propagate out to more architecturally granular regions at the external pole ^13^. These hypotheses are consistent with “core-periphery” ^32,85^ and “bow-tie” ^86^ descriptions of cortical networks.

The second gradient in the cerebral cortex distinguished ensembles associated with the representation of prediction and prediction error signals (anchored in nodes from the sensory and default mode networks) from those involved in the implementation of attentional modulation to set the precision of these signals (with nodes from frontoparietal and salience networks) ^36^, where “attention” is defined not in terms of properties of subjective awareness but as signals that modulate the firing rate of neurons. It has been hypothesized that the frontoparietal network estimates the precision of prediction signals or priors, possibly suppressing or inhibiting prediction ensembles whose priors are very low ^13^. In contrast, the salience network may alter the gain on prediction error signals as they propagate from the sensory periphery, reflecting confidence in the reliability and quality of incoming sensory information as well as its predicted relevance for allostasis and motor control ^13^. During some tasks, these modulatory networks cohere into a single “task positive” network ^87^ or a “multiple demand” system ^88^.

Our results lend support to the hypothesis that the cerebral cortex, the cerebellum, and the hippocampus all share common axes of functional organization. Regarding the cerebellum, we largely replicated across samples prior work on functional connectivity gradients in this structure ^38^. The most dominant gradient in the cerebellum has been characterized as a gradual transition from areas involved in motor function to those implicated in non-motor functions involved in cognitive, social, and emotional tasks ^38^. It is anchored at one end by the default mode and frontoparietal control networks and at the other end by the somatomotor and salience networks, consistent with the cortical model-error gradient. The second cerebellar gradient showed preferential functional connectivity with the default mode and somatomotor networks at one end and the frontoparietal and salience networks at the other ^89^. This gradient has been interpreted as reflective of differences in relation to “task-focus” ^38^ that preferentially engages networks in the presence of higher cognitive load ^87^; this is consistent with the cortical model-precision gradient in the present study.

Turning to the hippocampus, the most dominant gradient that has been identified in the hippocampus captures variation in functional connectivity along its longitudinal axis ^39–41^, consistent with evidence identifying gradual changes in anatomical connectivity, gene expression, and electrophysiological response properties along this axis ^68,90,91^. The largest difference in functional connectivity between the anterior and posterior subregions has been observed with the frontoparietal and salience networks, with the posterior (septal) hippocampus showing stronger positive connectivity with these networks, and the default mode and somatomotor networks at the anterior (uncal) end ^41^. Our results generally confirmed this pattern, which is consistent with the model-precision gradient of the cerebral cortex. The second hippocampal gradient seems to correspond to hippocampal microstructure, primarily isolating the subiculum from the CA subfields. This result is in line with prior evidence characterizing distinct subfields with variation in connectivity, computational roles, and myeloarchitectural maturation ^92–95^. Also consistent with our finding, the subiculum was found to show stronger functional connectivity with the default mode network than the other subfields, whereas CA1-3 showed stronger connectivity with the somatosensory, somatomotor, and visual networks ^41^; this gradient thus appears to be consistent with the model-error gradient in the cerebral cortex. These observations were confirmed with our vertex-wise analysis of correspondence in functional connectivity gradients across structures, suggesting that the model-error and model-precision gradients may represent the common axes of functional organization capturing the coordination of prediction, prediction error, and precision signals. This evidence has important implications for understanding the computational mechanisms underlying functional coordination across these structures.

### Coordination of cortical and cerebellar connectivity gradients

A traditional view of cortico-cerebellar interactions is that the cerebellum estimates the sensory state of the body by anticipating the consequences of a motor command ^9,20,42^, possibly as a means to compensate for the relatively slower temporal scale in which sensory feedback signals are processed in the cerebral cortex ^96,97^. In sensorimotor coordination, current evidence suggests that the cerebellar cortex (e.g., lobules V and VI) receives efferent copies of motor commands from Layer V of the primary motor cortex via the pontine nuclei ^98,99^ and predicts the expected sensory consequences of those commands ^100,101^. The inferior olive in the medulla oblongata is thought to play a critical role as one comparator of the predictions about the sensory consequences of motor commands from the cerebellum vs. the actual sensory input conveyed via afferent projections from the spinal cord ^7,102^. Prediction errors, i.e., the difference between the predicted and actual sensory inputs, are relayed to the premotor and primary motor cortices via the ventrolateral thalamus, as well as to motor neurons and interneurons in the lower brainstem and spinal cord to adjust motor movements online ^103^. These cerebellar-mediated sensory prediction error signals are important for refining motor outputs as well as future sensory predictions ^42^.

An evolutionary perspective on the cerebellum helps to elaborate and modify this view to hypothesize the functional value of the coordinated cortico-cerebellar connectivity gradients identified in this work. The brains of all major groups of vertebrates include a cerebellum ^104,105^. In fish, for example, the cerebellum-like structure allows the brain to model expected patterns of peripheral sensory input related to predictable water currents and the animal’s own movement, and adaptively filter the sensory consequences of these sensory signals, which in turn helps the fish detect unpredictable, behaviorally relevant sensory events and compute the corresponding sensory prediction errors more effectively ^106^. From this perspective, the cerebellum (and cerebellum-like structures) might be thought of as a sensory structure running a sensory model of the body in the world, which has been elaborated in land vertebrates, and particularly in mammals, in the service of complex motor control.

In humans, the cerebellum receives ascending sensory inputs from the dorsal spinocerebellar tract (via the medulla), while at the same time also receiving from the ventral spinocerebellar tract (via the pons) copies of motor commands originally sent to spinal motor neurons ^107^; these latter signals can be considered corollary discharge ^108^. In the cerebral cortex, corollary discharge of motor commands are thought to serve as sensory prediction signals ^13,15,109^ and the same may be true for corollary discharge signals reaching the cerebellum ^42,108,110^. Convergence of sensory predictions and incoming sense data would allow the cerebellum to compare the two sources of information, possibly resulting in the computation of sensory prediction errors. Both Purkinje cells and granule cells in the cerebellar cortex are thought to be involved in this comparison process ^107^. Given these lines of evidence, one possibility is that the cerebellum might send these modeled prediction error signals to the primary motor cortex to rapidly adjust motor control faster than cortical sensory prediction errors can be computed, in addition to sending its descending prediction signals to the effector organs via the brainstem. These cerebellar prediction errors might also be compared to the sensory prediction errors computed in the cerebral cortex.

This logic helps us speculate on the functional significance of cerebellar G_1_, whose connectivity patterns correspond to the model-error gradient within the cerebral cortex. For instance, one novel hypothesis is that, at one end, the cerebellum’s predictions about sensory prediction errors might be available to adjust motor and sensory predictions that originate in the default mode network, whose precision may be modulated by the frontoparietal network. These cerebellar prediction errors might be able to modulate activity of these networks faster than cortically computed sensory prediction errors, which are computed relatively more slowly. At the other end of this gradient, cerebellar modeled sensory prediction errors might be available to compare with sensory prediction errors modeled in the cerebral cortex, in the cortical sensory networks. Consistent with this hypothesis, current theories of cerebellar function posit that such predictive mechanisms may generalize to perceptual activity ^7,42,111–114^. For instance, in visual perception, the cerebellum is thought to be critical for predicting incoming sensory information based on sequence detection and updating predictions based on the statistics of the sensory environment ^115–120^. Visual information is conveyed polysynaptically from visual association cortical areas to cerebellar lobules VI and VII via the pons ^121,122^. From the perspective of the cerebellum, this information may represent sensory prediction error signals from the cerebral cortex. Visual inputs also reach the cerebellum from primary sensory receptors (e.g., via the superior colliculus) ^122^, which may allow the cerebellum to generate and update predictions about future sensory experiences. Collectively, one overarching hypothesis concerning cortico-cerebellar interaction is that both the cerebral cortex and the cerebellum are capable of computing sensory prediction errors and are possibly exchanging and comparing them to more efficiently update the brain’s internal model of its body in the world. This is just one example of a novel hypothesis suggested by the correspondence of connectivity gradients observed in this study, which provides a fruitful avenue for future research using anatomical, electrophysiological, and lesion data across species.

Recent evidence also suggests that the cerebellum may be involved in estimating the precision of sensory prediction errors, consistent with the organization of cerebellar G_2_ corresponding to the model-precision gradient in the cerebral cortex. During motor learning, the brain controls error sensitivity (i.e., the extent to which the brain changes the motor commands in the trial following an error) by learning relatively more from small and consistent errors than from larger and variable ones ^123,124^. This learning mechanism depends critically on the memory of errors that accumulates during training, which exists independently of two traditional forms of motor memory (memory of perturbations and of actions) ^123^. Although motor learning can occur on different time scales with different error sensitivities ^125^, the memory of errors is thought to exert its influence through the error sensitivity of the fast learning process ^123^. Therefore, one possibility is that the cerebellum rapidly estimates the reliability of sensory prediction errors, conveying this information to parts of the cerebral cortex (e.g., the premotor areas such as the anterior mid cingulate cortex within the salience network) where it is further used to update precision estimates about sensory prediction error signals.

Microanatomical and connectivity evidence further supports the hypothesis that the cerebellum can exert rapid modulation of signals in the cerebral cortex via coordinated gradients. The majority of cerebellar neurons are granule cells, which can generate action potentials that are relatively short-lived and at much higher frequencies than neurons in the cerebral cortex ^96^. Deep cerebellar nuclei, which are the gateway of cerebellar output, can also be modulated to fire up to 100+ Hz on average ^126^. These physiological properties may allow the cerebellum to rapidly modulate prediction, prediction error, and precision signals in the cerebral cortex in a domain-general fashion. Despite the fact that the cerebral cortex and the cerebellum are connected to each other only by way of polysynaptic circuits ^89,127^, numerous nonprimary sensorimotor (e.g., parietal association, parahippocampal, occipitotemporal, and prefrontal) areas of the cerebral cortex project to the cerebellar cortex via the cortico-ponto-cerebellar paths ^128^. The neocortical areas that project to specific parts of the cerebellar cortex via the pons are also the target of efferent projections from the same cerebellar cortical areas via the thalamus ^129–131^. These parallel, reciprocally-connected circuits might provide an anatomical substrate for the coordinated functional connectivity gradients identified in this study. Overall, our findings extend prior work in the functional parcellation of the cerebellum ^38,89,132–134^ by suggesting the potential mechanisms underlying the contribution of cortico-cerebellar interactions to the brain’s predictive processing.

### Coordination of cortical and hippocampal connectivity gradients

A traditional view of cortico-hippocampal interactions is that the cerebral cortex generates predictions based on the sensory statistics of the environment, whereas the hippocampus—which itself generates prediction signals—reweights and alters the cortical signals according to the goals of the organism ^43,44^. This mechanism draws upon the functional loop between the hippocampus, the entorhinal cortex, and the neocortex. Cortical afferents to the hippocampus carry highly compressed, multi-modal summaries of sensory information via the entorhinal cortex ^135^, whose Layer II and III project widely to the DG, CA1-CA4, and the subiculum via the perforant path ^91,136^. From the perspective of the hippocampus, these signals may represent prediction error signals from the cerebral cortex. Subcortical projections to the hippocampus include those from the medial septum, amygdala, anterior thalamic nuclei, supramammillary nucleus of the hypothalamus, as well as several brainstem nuclei such as the ventral tegmental area, periaqueductal gray, and locus coeruleus ^136,137^, possibly carrying information about the sensory state of the body including energy and metabolic requirements. The hippocampus may integrate information coming from these sources in its own internal architecture to generate predictions about future experiences; this mechanism may as well be one way in which the hippocampus performs the reweighting and adjustment of cortical predictions.

Hippocampal signals reach back out to the cerebral cortex through various routes to achieve these adjustments. Specifically, CA1 and the subiculum in turn project out to Layer V and VI of the entorhinal cortex ^135^ as well as widespread multimodal association areas in the cerebral cortex including the medial frontal cortex, temporal pole, orbitofrontal cortex, anterior and posterior cingulate cortices, parietal and inferotemporal cortices ^138,139^ and to some extent lateral frontal cortex ^138^. From the perspective of the cerebral cortex, these signals may represent prediction errors consisting of multi-modal information generated from within the hippocampus ^140–142^, which could then be unpacked and particularized as they are integrated with the cerebral cortex’s internal model. By adjusting the representations in the cerebral cortex, the hippocampus may help ensure that the subsequent prediction signals generated based on the cortex’s internal model are not slaves to the statistics of the external sensory environment and instead more in line with the goals of the animal (i.e., weighted for the current and perdicted conditions of the body’s internal environment) ^44^.

One novel hypothesis emerging from the findings of the current study is that the hippocampal connectivity gradients together characterize the role of the hippocampus in adjusting prediction signals in the service of allostasis along the entire cortical sheet. That is, the two most dominant connectivity gradients of the hippocampus might actually be considered a single gradient with three functional anchors along the longitudinal axis. At the posterior end of the hippocampus (corresponding to the bottom 10% of hippocampal G_1_ in the current study), functional connectivity was stronger with the cortical attentional networks, suggesting the role of hippocampal neurons in this area in tuning the precision signals acting on prediction and prediction error signals in the cerebral cortex. The posterior (septal in rodents) hippocampus receives ascending interoceptive prediction errors from the medial septum ^143^ and from the supramammilary nucleus of the hypothalamus ^144^, which could reweight sensory statistics based on the internal state of the body. The medial septum is critical for the generation of theta frequency oscillations observed in the hippocampus ^145,146^, which are important for the hippocampus’ role in generating predicted sequences of sensory events ^43^. The medial septum also mediates the effect of vagus nerve (parasympathetic afferents) stimulation on hippocampal theta oscillations ^147–149^. This may suggest the posterior hippocampus’ involvement in using interoceptive information to guide processing of event sequences.

In the middle of the hippocampus (corresponding to the top 10% of hippocampal G_1_ and G_2_), functional connectivity was stronger with the default mode network in the cerebral cortex. This network is critical for initiating prediction signals constituting the cerebral cortex’s internal model constructed from past experiences ^13,150^. This network is also key for conceptual processing ^151^, and on one hypothesis prediction signals can be thought of as low-dimensional, conceptual representations that guide the conceptualization (i.e., categorization) of incoming sensory information in the service of efficient bodily regulation ^13^. The middle portion of the hippocampus, therefore, may interact with the default mode network in the cerebral cortex to categorize sensory inputs and give them meaning, where “meaning” includes the generation of visceromotor and motor action plans to deal with that particular event in a specific context ^13^. In this way, the hippocampus may also contribute to interoceptive predictions – by sending anticipated sensory consequences of visceromotor changes to the primary interoceptive cortex ^15,17^ and therefore a change in affect (e.g., mood).

At the anterior end of the hippocampus (corresponding to the bottom 10% of hippocampal G_2_), functional connectivity was stronger with the somatosensory and somatomotor cortices, suggesting the role of hippocampal neurons in this area in sending motor and sensory predictions to the cerebral cortex, possibly along with visceromotor predictions given the presence of visceral maps within the primary motor cortex ^152,153^. Preferential connectivity with the sensorimotor cortices in the anterior (vs. posterior) hippocampus in humans is consistent with available evidence ^40,68,154^. In rodents, the temporal (i.e., anterior) two thirds of CA1 passing through the longitudinal association bundles project to the primary visceral sensory area and the supplementary somatosensory area ^155^, which may correspond to the functional connection observed in this study. The attentional-conceptual-(viscero)sensorimotor gradient in the hippocampus, therefore, may characterize the hippocampus’ contribution to predictive processing in the brain, which involves the refinement of representations in the cerebral cortex regardless of whether they are content-based or modulatory. Such tripartite organization of hippocampal function is consistent with prior work in non-human primates and rodents characterizing the topographic organization of hippocampal-entorhinal interconnections, where the anterior, middle, and posterior subregions of the hippocampus exhibit preferential connections with the medial, intermediate, and lateral bands of the entorhinal cortex, respectively ^156–158^. This entorhinal mediolateral gradient in turn appears to be associated with distinct patterns of functional connectivity in humans, although evidence is still preliminary given the limited spatial coverage of the data ^159^. Future work examining whole-brain high-resolution fMRI data is warranted to clarify the mechanisms underlying signal flow and integration along a neocortical, entorhinal, and hippocampal functional loop.

Notably, our findings meaningfully extend the existing accounts of the functional organization of hippocampus identified in humans ^160–163^, by demonstrating evidence supporting the role of cortico-hippocampal interactions in domain-general computational processes. Much of the prior work examining longitudinal-axis functional specialization within the hippocampus has focused on characterizing it by distinct patterns of functional interaction with nearby structures in the medial temporal lobe or with a broader set of cortical regions that are canonically considered part of the default mode network; this specialization has been most typically linked to different aspects of episodic memory ^154,161,164–171^. Our findings are consistent with recent evidence showing that hippocampal functional specialization along its longitudinal axis reflects its relevance not just for memory but across multiple functional domains ^172^, thus underscoring the importance of adopting a more domain-general view of hippocampal function than traditionally thought in the memory literature.

### Coordination of cerebellar and hippocampal connectivity gradients

Finally, the current findings of coordinated connectivity gradients offer novel insights to probe observations that are relatively understudied in the literature, such as the interaction between the cerebellum and the hippocampus. Emerging evidence suggests the existence of a cerebello-hippocampal learning system ^45–49,173,174^, although its computational and functional architecture are relatively less well studied when compared to the other learning systems discussed. Viral tracing studies have so far identified polysynaptic connections between these structures mediated by regions including the supramammillary nucleus of the hypothalamus, medial septum, and ventrolateral/laterodorsal thalamus ^48,175^. There is also evidence pointing to the existence of direct connections between cerebellar and hippocampal subregions in humans ^176^. The present findings reinforce the importance of testing specific hypotheses, for instance, about event segmentation and sequence processing in which both structures have been (separately) implicated ^43,117,177–180^. Future work should investigate the complementary contributions of cortico-cerebellar, cortico-hippocampal, and cerebello-hippocampal interactions to the brain’s internal model, which might be characterized by their dissociable involvement in processing different types of information and/or on different timescales.

### Conclusions

The present results offer the opportunity to synthesize evidence across literatures, each targeting a different set of brain regions, into a common neurocomputational framework based on the principles of predictive processing. Our hypotheses, while speculative, illustrate the value of connectivity gradients in innovating specific questions about the computational aspects of brain function, with the model-error and model-precision gradients as two common axes of information processing in the brain. Future work might specifically address these questions, as well as probe modulation of connectivity gradient coordination across structures by explicit task demands or by clinical conditions in which neural mechanisms subserving predictive processing are hypothesized to be dysfunctional ^181,182^. The results might offer a coherent, neurobiologically-inspired research program to unite the study of mind and behavior, collapsing the artificial boundaries between cognitive, perceptual, affective, motor, and even social phenomena. This evidence might also provide a common framework for understanding more broadly the neurocomputational basis of mental disorders, neurodegenerative disorders, and physical disorders.

## Methods

A full description of the datasets, data processing, and analytical approaches is provided as part of **Supplementary Methods**. Briefly, we analyzed two large resting-state fMRI datasets, both of which are publicly available. The primary dataset consisted of 1,003 participants from the HCP WU-Minn Consortium ^52^ (HCP1200 2017 data release; *M*_age_ = 28.71, *SD*_age_ = 3.71, 470 males, 533 females; four 15 min runs per participant). Specifically, we utilized the group average preprocessed dense functional connectome data generated via the HCP pipelines ^183,184^. These data took the form of a 91,282 × 91,282 matrix representing the magnitude of functional connectivity between all cortical vertices and subcortical voxels; the hippocampus was represented as part of the subcortical volume space ^183^. A secondary, validation dataset was defined with an independent group of 1,139 participants from the GSP (*M*_age_ = 21.24, *SD*_age_ = 2.70, 467 males, 672 females; two 6 min runs per participant) ^53,54^. We performed preprocessing of the GSP dataset using a surface-based pipeline ^58,185^, after which we generated the group average whole-brain vertex-/voxel-wise functional connectivity matrix.

From these group-level functional connectivity matrices, we extracted (1) cortico-cortical, (2) cerebello-cortical, and (3) hippocampo-cortical connections, which were used as input for diffusion map embedding, a non-linear data dimensionality reduction technique that allows calculation of functional connectivity gradients as a low-dimensional representation of spatial variation in connectivity profiles ^50,51^. We performed post hoc characterization and interpretation of the observed functional gradients at various levels, including comparisons with the topography of canonical functional networks ^54^ and examination of gradient value distributions across major hippocampal subfields ^62^. To interpret the correspondence between cerebellar/hippocampal and cerebral cortical connectivity gradients, we computed gradient-weighted functional connectivity in cortical space ^36,59^ and quantified the degree of spatial correlation while non-parametrically accounting for autocorrelations ^63^. Finally, to further demonstrate the correspondence between the cortical, cerebellar, and hippocampal functional gradients in terms of connectivity, we performed a seed-based functional connectivity analysis by targeting the vertices/voxels that occupied the top and bottom 10% of each gradient. This procedure enabled triangulation of intrinsic functional connectivity between distinct subregions of the three structures that corresponded to a common connectivity gradient.

## Data Availability

The HCP dataset is publicly available at https://db.humanconnectome.org. The GSP dataset is publicly available at doi: https://dataverse.harvard.edu/dataset.xhtml?persistentId=10.7910/DVN/25833. The fMRI data derivatives generated in this work will be made available at https://github.com/yutakatsumi/PredProcGradients.

## Acknowledgements

This work was supported by the NSF (BCS 1947972 to L.F.B.) and the NIH (R01 MH113234 to L.F.B. and B.C.D., R01 MH109464 to L.F.B., and U01 CA193632 to L.F.B.).

## References

1. Friston, K., FitzGerald, T., Rigoli, F., Schwartenbeck, P. & Pezzulo, G. Active Inference: A Process Theory. Neural Comput. 29, 1–49 (2017).

2. Hutchinson, J. B. & Barrett, L. F. The Power of Predictions: An Emerging Paradigm for Psychological Research. Curr. Dir. Psychol. Sci. 28, 280–291 (2019).

3. Berkes, P., Orbán, G., Lengyel, M. & Fiser, J. Spontaneous Cortical Activity Reveals Hallmarks of an Optimal Internal Model of the Environment. Science 331, 83–87 (2011).

4. Friston, K. The free-energy principle: a unified brain theory? Nat. Rev. Neurosci. 11, 127–138 (2010).

5. Jordan, R. & Keller, G. B. Opposing Influence of Top-down and Bottom-up Input on Excitatory Layer 2/3 Neurons in Mouse Primary Visual Cortex. Neuron 108, 1194–1206.e5 (2020).

6. Rao, R. P. N. & Ballard, D. H. Predictive coding in the visual cortex: a functional interpretation of some extra-classical receptive-field effects. Nat. Neurosci. 2, 79–87 (1999).

7. Ito, M. Control of mental activities by internal models in the cerebellum. Nat. Rev. Neurosci. 9, 304–313 (2008).

8. Kawato, M. Internal models for motor control and trajectory planning. Curr. Opin. Neurobiol. 9, 718–727 (1999).

9. Wolpert, D. M., Miall, R. C. & Kawato, M. Internal models in the cerebellum. Trends Cogn. Sci. 2, 338–347 (1998).

10. Lamme, V. A. F. & Roelfsema, P. R. The distinct modes of vision offered by feedforward and recurrent processing. Trends Neurosci. 23, 571–579 (2000).

11. Feldman, H. & Friston, K. J. Attention, uncertainty, and free-energy. Front. Hum. Neurosci. 4, 215 (2010).

12. Kanai, R., Komura, Y., Shipp, S. & Friston, K. Cerebral hierarchies: predictive processing, precision and the pulvinar. Philos. Trans. R. Soc. B Biol. Sci. 370, 20140169 (2015).

13. Barrett, L. F. The theory of constructed emotion: an active inference account of interoception and categorization. Soc. Cogn. Affect. Neurosci. 12, 1–23 (2017).

14. Parr, T. & Friston, K. J. Attention or salience? Curr. Opin. Psychol. 29, 1–5 (2019).

15. Barrett, L. F. & Simmons, W. K. Interoceptive predictions in the brain. Nat. Rev. Neurosci. 16, 419–429 (2015).

16. Chanes, L. & Barrett, L. F. Redefining the Role of Limbic Areas in Cortical Processing. Trends Cogn. Sci. 20, 96–106 (2016).

17. Kleckner, I. R. et al. Evidence for a large-scale brain system supporting allostasis and interoception in humans. Nat. Hum. Behav. 1, 1–14 (2017).

18. Keller, G. B. & Mrsic-Flogel, T. D. Predictive Processing: A Canonical Cortical Computation. Neuron 100, 424–435 (2018).

19. Picard, F. & Friston, K. Predictions, perception, and a sense of self. Neurology 83, 1112–1118 (2014).

20. Shadmehr, R., Smith, M. A. & Krakauer, J. W. Error Correction, Sensory Prediction, and Adaptation in Motor Control. Annu. Rev. Neurosci. 33, 89–108 (2010).

21. Barron, H. C., Auksztulewicz, R. & Friston, K. Prediction and memory: A predictive coding account. Prog. Neurobiol. 192, 101821 (2020).

22. Gravina, M. T. & Sederberg, P. B. The neural architecture of prediction over a continuum of spatiotemporal scales. Curr. Opin. Behav. Sci. 17, 194–202 (2017).

23. Liu, K., Sibille, J. & Dragoi, G. Generative Predictive Codes by Multiplexed Hippocampal Neuronal Tuplets. Neuron 99, 1329–1341.e6 (2018).

24. Sherman, B. E. & Turk-Browne, N. B. Statistical prediction of the future impairs episodic encoding of the present. Proc. Natl. Acad. Sci. 117, 22760–22770 (2020).

25. Hong, S.-J. et al. Toward a connectivity gradient-based framework for reproducible biomarker discovery. NeuroImage 223, 117322 (2020).

26. Huntenburg, J. M., Bazin, P.-L. & Margulies, D. S. Large-Scale Gradients in Human Cortical Organization. Trends Cogn. Sci. 22, 21–31 (2018).

27. Katsumi, Y., Quigley, K. & Barrett, L. F. Situating allostasis and interoception at the core of human brain function. PsyArxiv (2021) doi:10.31234/osf.io/wezv8.

28. Biswal, B., Zerrin Yetkin, F., Haughton, V. M. & Hyde, J. S. Functional connectivity in the motor cortex of resting human brain using echo-planar mri. Magn. Reson. Med. 34, 537–541 (1995).

29. Buckner, R. L., Krienen, F. M. & Yeo, B. T. T. Opportunities and limitations of intrinsic functional connectivity MRI. Nat. Neurosci. 16, 832–837 (2013).

30. Bethlehem, R. A. I. et al. Dispersion of functional gradients across the adult lifespan. NeuroImage 222, 117299 (2020).

31. Hong, S.-J. et al. Atypical functional connectome hierarchy in autism. Nat. Commun. 10, 1–13 (2019).

32. Margulies, D. S. et al. Situating the default-mode network along a principal gradient of macroscale cortical organization. Proc. Natl. Acad. Sci. 113, 12574–12579 (2016).

33. Paquola, C. et al. Microstructural and functional gradients are increasingly dissociated in transmodal cortices. PLOS Biol. 17, e3000284 (2019).

34. Raut, R. V., Snyder, A. Z. & Raichle, M. E. Hierarchical dynamics as a macroscopic organizing principle of the human brain. Proc. Natl. Acad. Sci. 117, 20890–20897 (2020).

35. Shafiei, G. et al. Topographic gradients of intrinsic dynamics across neocortex. http://biorxiv.org/lookup/doi/10.1101/2020.07.03.186916 (2020) xdoi:10.1101/2020.07.03.186916.

36. Zhang, J. et al. Intrinsic functional connectivity is organized as three interdependent gradients. Sci. Rep. 9, 1–14 (2019).

37. Dong, D. et al. Compression of Cerebellar Functional Gradients in Schizophrenia. Schizophr. Bull. (2020) doi:10.1093/schbul/sbaa016.

38. Guell, X., Schmahmann, J. D., Gabrieli, J. D. & Ghosh, S. S. Functional gradients of the cerebellum. eLife 7, e36652 (2018).

39. Kharabian Masouleh, S., Plachti, A., Hoffstaedter, F., Eickhoff, S. & Genon, S. Characterizing the gradients of structural covariance in the human hippocampus. NeuroImage 218, 116972 (2020).

40. Przeździk, I., Faber, M., Fernández, G., Beckmann, C. F. & Haak, K. V. The functional organisation of the hippocampus along its long axis is gradual and predicts recollection. Cortex 119, 324–335 (2019).

41. Vos de Wael, R. et al. Anatomical and microstructural determinants of hippocampal subfield functional connectome embedding. Proc. Natl. Acad. Sci. 115, 10154–10159 (2018).

42. Sokolov, A. A., Miall, R. C. & Ivry, R. B. The Cerebellum: Adaptive Prediction for Movement and Cognition. Trends Cogn. Sci. 21, 313–332 (2017).

43. Buzsáki, G. & Tingley, D. Space and Time: The Hippocampus as a Sequence Generator. Trends Cogn. Sci. 22, 853–869 (2018).

44. Kumaran, D., Hassabis, D. & McClelland, J. L. What Learning Systems do Intelligent Agents Need? Complementary Learning Systems Theory Updated. Trends Cogn. Sci. 20, 512–534 (2016).

45. Babayan, B. M. et al. A hippocampo-cerebellar centred network for the learning and execution of sequence-based navigation. Sci. Rep. 7, 17812 (2017).

46. Iglói, K. et al. Interaction Between Hippocampus and Cerebellum Crus I in Sequence-Based but not Place-Based Navigation. Cereb. Cortex 25, 4146–4154 (2015).

47. Onuki, Y., Van Someren, E. J. W., De Zeeuw, C. I. & Van der Werf, Y. D. Hippocampal–Cerebellar Interaction During Spatio-Temporal Prediction. Cereb. Cortex 25, 313–321 (2015).

48. Watson, T. C. et al. Anatomical and physiological foundations of cerebello-hippocampal interaction. eLife 8, e41896 (2019).

49. Yu, W. & Krook-Magnuson, E. Cognitive Collaborations: Bidirectional Functional Connectivity Between the Cerebellum and the Hippocampus. Front. Syst. Neurosci. 9, (2015).

50. Coifman, R. R. & Lafon, S. Diffusion maps. Appl. Comput. Harmon. Anal. 21, 5–30 (2006).

51. Vos de Wael, R. et al. BrainSpace: a toolbox for the analysis of macroscale gradients in neuroimaging and connectomics datasets. Commun. Biol. 3, 1–10 (2020).

52. Van Essen, D. C. et al. The WU-Minn Human Connectome Project: An overview. NeuroImage 80, 62–79 (2013).

53. Holmes, A. J. et al. Brain Genomics Superstruct Project initial data release with structural, functional, and behavioral measures. Sci. Data 2, 1–16 (2015).

54. Yeo, B. T. T. et al. The organization of the human cerebral cortex estimated by intrinsic functional connectivity. J. Neurophysiol. 106, 1125–1165 (2011).

55. Barbas, H. General Cortical and Special Prefrontal Connections: Principles from Structure to Function. Annu. Rev. Neurosci. 38, 269–289 (2015).

56. García-Cabezas, M. Á., Zikopoulos, B. & Barbas, H. The Structural Model: a theory linking connections, plasticity, pathology, development and evolution of the cerebral cortex. Brain Struct. Funct. 224, 985–1008 (2019).

57. Beul, S. F. & Hilgetag, C. C. Towards a “canonical” agranular cortical microcircuit. Front. Neuroanat. 8, (2015).

58. Kong, R. et al. Spatial Topography of Individual-Specific Cortical Networks Predicts Human Cognition, Personality, and Emotion. Cereb. Cortex 29, 2533–2551 (2019).

59. Guell, X. et al. Functional Territories of Human Dentate Nucleus. Cereb. Cortex (2020) doi:10.1093/cercor/bhz247.

60. Li, Q. et al. Atypical neural topographies underpin dysfunctional pattern separation in temporal lobe epilepsy. Brain (2021) doi:10.1093/brain/awab121.

61. Yang, S. et al. The thalamic functional gradient and its relationship to structural basis and cognitive relevance. NeuroImage 218, 116960 (2020).

62. Kulaga-Yoskovitz, J. et al. Multi-contrast submillimetric 3 Tesla hippocampal subfield segmentation protocol and dataset. Sci. Data 2, 1–9 (2015).

63. Alexander-Bloch, A. F. et al. On testing for spatial correspondence between maps of human brain structure and function. NeuroImage 178, 540–551 (2018).

64. Blazquez Freches, G. et al. Principles of temporal association cortex organisation as revealed by connectivity gradients. Brain Struct. Funct. 225, 1245–1260 (2020).

65. Faber, M., Przezdzik, I., Fernandez, G., Haak, K. V. & Beckmann, C. F. Overlapping connectivity gradients in the anterior temporal lobe underlie semantic cognition. bioRxiv 2020.05.28.121137 (2020) doi:10.1101/2020.05.28.121137.

66. Haak, K. V., Marquand, A. F. & Beckmann, C. F. Connectopic mapping with resting-state fMRI. NeuroImage 170, 83–94 (2018).

67. Paquola, C. et al. Convergence of cortical types and functional motifs in the mesiotemporal lobe. bioRxiv 2020.06.12.148643 (2020) doi:10.1101/2020.06.12.148643.

68. Vogel, J. W. et al. A molecular gradient along the longitudinal axis of the human hippocampus informs large-scale behavioral systems. Nat. Commun. 11, 1–17 (2020).

69. Haueis, P. Multiscale modeling of cortical gradients: the role of mesoscale circuits for linking macro-and microscale gradients of cortical organization and hierarchical information processing. NeuroImage 117846 (2021) doi:10.1016/j.neuroimage.2021.117846.

70. Zhang, J. et al. Topography Impacts Topology: Anatomically Central Areas Exhibit a “High-Level Connector” Profile in the Human Cortex. Cereb. Cortex 30, 1357–1365 (2020).

71. Marquand, A. F., Haak, K. V. & Beckmann, C. F. Functional corticostriatal connection topographies predict goal-directed behaviour in humans. Nat. Hum. Behav. 1, 1–9 (2017).

72. Mesulam, M. From sensation to cognition. Brain 121, 1013–1052 (1998).

73. Finlay, B. L. & Uchiyama, R. Developmental mechanisms channeling cortical evolution. Trends Neurosci. 38, 69–76 (2015).

74. Barbas, H. & García-Cabezas, M. Á. Motor cortex layer 4: less is more. Trends Neurosci. 38, 259–261 (2015).

75. Zhang, S. et al. Long-Range and Local Circuits for Top-Down Modulation of Visual Cortical Processing. Science 345, 660–665 (2014).

76. Leinweber, M., Ward, D. R., Sobczak, J. M., Attinger, A. & Keller, G. B. A Sensorimotor Circuit in Mouse Cortex for Visual Flow Predictions. Neuron 95, 1420–1432.e5 (2017).

77. Fiser, J., Chiu, C. & Weliky, M. Small modulation of ongoing cortical dynamics by sensory input during natural vision. Nature 431, 573–578 (2004).

78. Gallant, J. L., Connor, C. E. & Van Essen, D. C. Neural activity in areas V1, V2 and V4 during free viewing of natural scenes compared to controlled viewing. NeuroReport 9, 85–89 (1998).

79. Keller, G. B., Bonhoeffer, T. & Hübener, M. Sensorimotor Mismatch Signals in Primary Visual Cortex of the Behaving Mouse. Neuron 74, 809–815 (2012).

80. Livingstone, M. S., Freeman, D. C. & Hubel, D. H. Visual responses in V1 of freely viewing monkeys. Cold Spring Harb. Symp. Quant. Biol. 61, 27–37 (1996).

81. Vinje, W. E. & Gallant, J. L. Sparse Coding and Decorrelation in Primary Visual Cortex During Natural Vision. Science 287, 1273–1276 (2000).

82. Sillito, A. M. & Jones, H. E. Corticothalamic interactions in the transfer of visual information. Philos. Trans. R. Soc. Lond. B. Biol. Sci. 357, 1739–1752 (2002).

83. Huntenburg, J. M. et al. A Systematic Relationship Between Functional Connectivity and Intracortical Myelin in the Human Cerebral Cortex. Cereb. Cortex 27, 981–997 (2017).

84. Pezzulo, G., Zorzi, M. & Corbetta, M. The secret life of predictive brains: what’s spontaneous activity for? PsyArxiv (2020) doi:10.31234/osf.io/qus3h.

85. Principles of behavioral and cognitive neurology. (Oxford University Press, 2000).

86. Markov, N. T. et al. Cortical High-Density Counterstream Architectures. Science 342, (2013).

87. Fox, M. D. et al. The human brain is intrinsically organized into dynamic, anticorrelated functional networks. Proc. Natl. Acad. Sci. 102, 9673–9678 (2005).

88. Assem, M., Glasser, M. F., Van Essen, D. C. & Duncan, J. A Domain-General Cognitive Core Defined in Multimodally Parcellated Human Cortex. Cereb. Cortex 30, 4361–4380 (2020).

89. Buckner, R. L., Krienen, F. M., Castellanos, A., Diaz, J. C. & Yeo, B. T. T. The organization of the human cerebellum estimated by intrinsic functional connectivity. J. Neurophysiol. 106, 2322–2345 (2011).

90. Fanselow, M. S. & Dong, H.-W. Are the Dorsal and Ventral Hippocampus Functionally Distinct Structures? Neuron 65, 7–19 (2010).

91. Strange, B. A., Witter, M. P., Lein, E. S. & Moser, E. I. Functional organization of the hippocampal longitudinal axis. Nat. Rev. Neurosci. 15, 655–669 (2014).

92. Ábrahám, H. et al. Myelination in the human hippocampal formation from midgestation to adulthood. Int. J. Dev. Neurosci. 28, 401–410 (2010).

93. Aggleton, J. P. Multiple anatomical systems embedded within the primate medial temporal lobe: Implications for hippocampal function. Neurosci. Biobehav. Rev. 36, 1579–1596 (2012).

94. Rolls, E. T. Pattern separation, completion, and categorisation in the hippocampus and neocortex. Neurobiol. Learn. Mem. 129, 4–28 (2016).

95. van Strien, N. M., Cappaert, N. L. M. & Witter, M. P. The anatomy of memory: an interactive overview of the parahippocampal–hippocampal network. Nat. Rev. Neurosci. 10, 272–282 (2009).

96. Sultan, F. et al. Unravelling cerebellar pathways with high temporal precision targeting motor and extensive sensory and parietal networks. Nat. Commun. 3, 924 (2012).

97. Tanaka, H., Ishikawa, T., Lee, J. & Kakei, S. The Cerebro-Cerebellum as a Locus of Forward Model: A Review. Front. Syst. Neurosci. 14, (2020).

98. Ishikawa, T., Tomatsu, S., Izawa, J. & Kakei, S. The cerebro-cerebellum: Could it be loci of forward models? Neurosci. Res. 104, 72–79 (2016).

99. Tomatsu, S. et al. Information processing in the hemisphere of the cerebellar cortex for control of wrist movement. J. Neurophysiol. 115, 255–270 (2015).

100. Herzfeld, D. J., Kojima, Y., Soetedjo, R. & Shadmehr, R. Encoding of action by the Purkinje cells of the cerebellum. Nature 526, 439–442 (2015).

101. Laurens, J., Meng, H. & Angelaki, D. E. Computation of linear acceleration through an internal model in the macaque cerebellum. Nat. Neurosci. 16, 1701–1708 (2013).

102. De Zeeuw, C. I. et al. Microcircuitry and function of the inferior olive. Trends Neurosci. 21, 391–400 (1998).

103. Ashida, R., Cerminara, N. L., Brooks, J. & Apps, R. Principles of organization of the human cerebellum: macro-and microanatomy. in Handbook of Clinical Neurology vol. 154 45–58 (Elsevier, 2018).

104. Montgomery, J. C., Bodznick, D. & Yopak, K. E. The Cerebellum and Cerebellum-Like Structures of Cartilaginous Fishes. Brain. Behav. Evol. 80, 152–165 (2012).

105. Voogd, J. & Glickstein, M. The anatomy of the cerebellum. Trends Neurosci. 21, 370–375 (1998).

106. Bratby, P., Montgomery, J. & Sneyd, J. A biophysical model of adaptive noise filtering in the shark brain. Bull. Math. Biol. 76, 455–475 (2014).

107. Bastian, A. J. & Lisberger, S. G. The Cerebellum. in Principles of Neural Science (McGraw Hill, 2021).

108. Straka, H., Simmers, J. & Chagnaud, B. P. A New Perspective on Predictive Motor Signaling. Curr. Biol. 28, R232–R243 (2018).

109. Adams, R. A., Shipp, S. & Friston, K. J. Predictions not commands: active inference in the motor system. Brain Struct. Funct. 218, 611–643 (2013).

110. Person, A. L. Corollary Discharge Signals in the Cerebellum. Biol. Psychiatry Cogn. Neurosci. Neuroimaging 4, 813–819 (2019).

111. Adamaszek, M. et al. Consensus Paper: Cerebellum and Emotion. The Cerebellum 16, 552–576 (2017).

112. Bareš, M. et al. Consensus paper: Decoding the Contributions of the Cerebellum as a Time Machine. From Neurons to Clinical Applications. The Cerebellum 18, 266–286 (2019).

113. Baumann, O. et al. Consensus Paper: The Role of the Cerebellum in Perceptual Processes. The Cerebellum 14, 197–220 (2015).

114. Koziol, L. F. et al. Consensus Paper: The Cerebellum’s Role in Movement and Cognition. The Cerebellum 13, 151–177 (2014).

115. Braitenberg, V., Heck, D. & Sultan, F. The detection and generation of sequences as a key to cerebellar function: Experiments and theory. Behav. Brain Sci. 20, 229–245 (1997).

116. Molinari, M. Cerebellum and procedural learning: evidence from focal cerebellar lesions. Brain 120, 1753–1762 (1997).

117. Molinari, M. et al. Cerebellum and Detection of Sequences, from Perception to Cognition. The Cerebellum 7, 611–615 (2008).

118. Molinari, M. & Masciullo, M. The Implementation of Predictions During Sequencing. Front. Cell. Neurosci. 13, (2019).

119. Roth, M. J., Synofzik, M. & Lindner, A. The Cerebellum Optimizes Perceptual Predictions about External Sensory Events. Curr. Biol. 23, 930–935 (2013).

120. Synofzik, M., Lindner, A. & Thier, P. The Cerebellum Updates Predictions about the Visual Consequences of One’s Behavior. Curr. Biol. 18, 814–818 (2008).

121. Bjaalie, J. G. & Brodal, P. Visual pathways to the cerebellum: Segregation in the pontine nuclei of terminal fields from different visual cortical areas in the cat. Neuroscience 29, 95–107 (1989).

122. Stein, J. F. & Glickstein, M. Role of the cerebellum in visual guidance of movement. Physiol. Rev. 72, 967–1017 (1992).

123. Herzfeld, D. J., Vaswani, P. A., Marko, M. K. & Shadmehr, R. A memory of errors in sensorimotor learning. Science 6 (2014).

124. Wei, K. & Körding, K. Relevance of Error: What Drives Motor Adaptation? J. Neurophysiol. 101, 655–664 (2009).

125. Smith, M. A., Ghazizadeh, A. & Shadmehr, R. Interacting Adaptive Processes with Different Timescales Underlie Short-Term Motor Learning. PLOS Biol. 4, e179 (2006).

126. Thach, W. T. Discharge of cerebellar neurons related to two maintained postures and two prompt movements. I. Nuclear cell output. J. Neurophysiol. 33, 527–536 (1970).

127. Schmahmann, J. D. & Pandya, D. N. The Cerebrocerebellar System. in International Review of Neurobiology (ed. Schmahmann, J. D.) vol. 41 31–60 (Academic Press, 1997).

128. Apps, R. & Watson, T. C. Cerebro-Cerebellar Connections. in Handbook of the Cerebellum and Cerebellar Disorders (eds. Manto, M., Schmahmann, J. D., Rossi, F., Gruol, D. L. & Koibuchi, N.) 1131–1153 (Springer Netherlands, 2013). doi:10.1007/978-94-007-1333-8_48.

129. Glickstein, M., May, J. G. & Mercier, B. E. Corticopontine projection in the macaque: The distribution of labelled cortical cells after large injections of horseradish peroxidase in the pontine nuclei. J. Comp. Neurol. 235, 343–359 (1985).

130. Kelly, R. M. & Strick, P. L. Cerebellar Loops with Motor Cortex and Prefrontal Cortex of a Nonhuman Primate. J. Neurosci. 23, 8432–8444 (2003).

131. Schmahmann, J. D. From movement to thought: Anatomic substrates of the cerebellar contribution to cognitive processing. Hum. Brain Mapp. 4, 174–198 (1996).

132. Guell, X., Gabrieli, J. D. E. & Schmahmann, J. D. Triple representation of language, working memory, social and emotion processing in the cerebellum: convergent evidence from task and seed-based resting-state fMRI analyses in a single large cohort. NeuroImage 172, 437–449 (2018).

133. Ji, J. L. et al. Mapping the human brain’s cortical-subcortical functional network organization. NeuroImage 185, 35–57 (2019).

134. King, M., Hernandez-Castillo, C. R., Poldrack, R. A., Ivry, R. B. & Diedrichsen, J. Functional boundaries in the human cerebellum revealed by a multi-domain task battery. Nat. Neurosci. 22, 1371–1378 (2019).

135. Witter, M. P., Doan, T. P., Jacobsen, B., Nilssen, E. S. & Ohara, S. Architecture of the Entorhinal Cortex A Review of Entorhinal Anatomy in Rodents with Some Comparative Notes. Front. Syst. Neurosci. 11, 46 (2017).

136. Insausti, R. & Amaral, D. G. Hippocampal Formation. in The Human Nervous System 896–942 (Elsevier, 2012). doi:10.1016/B978-0-12-374236-0.10024-0.

137. Amaral, D. G. & Cowan, W. M. Subcortical afferents to the hippocampal formation in the monkey. J. Comp. Neurol. 189, 573–591 (1980).

138. Barbas, H. & Blatt, G. J. Topographically specific hippocampal projections target functionally distinct prefrontal areas in the rhesus monkey. Hippocampus 5, 511–533 (1995).

139. Insausti, R. & Muñoz, M. Cortical projections of the non-entorhinal hippocampal formation in the cynomolgus monkey (Macaca fascicularis). Eur. J. Neurosci. 14, 435–451 (2001).

140. Duncan, K., Ketz, N., Inati, S. J. & Davachi, L. Evidence for area CA1 as a match/mismatch detector: A high-resolution fMRI study of the human hippocampus. Hippocampus 22, 389–398 (2012).

141. Kumaran, D. & Maguire, E. A. Which computational mechanisms operate in the hippocampus during novelty detection? Hippocampus 17, 735–748 (2007).

142. Lisman, J. E. & Grace, A. A. The Hippocampal-VTA Loop: Controlling the Entry of Information into Long-Term Memory. Neuron 46, 703–713 (2005).

143. Suarez, A. N. et al. Gut vagal sensory signaling regulates hippocampus function through multi-order pathways. Nat. Commun. 9, 1–15 (2018).

144. Vertes, R. P. Chapter 7 - Major diencephalic inputs to the hippocampus: supramammillary nucleus and nucleus reuniens. Circuitry and function. in Progress in Brain Research (eds. O’Mara, S. & Tsanov, M.) vol. 219 121–144 (Elsevier, 2015).

145. Tsanov, M. Differential and complementary roles of medial and lateral septum in the orchestration of limbic oscillations and signal integration. Eur. J. Neurosci. 48, 2783–2794 (2018).

146. Colgin, L. L. Mechanisms and Functions of Theta Rhythms. Annu. Rev. Neurosci. 36, 295–312 (2013).

147. Broncel, A., Bocian, R., Kłos-Wojtczak, P. & Konopacki, J. Vagus nerve stimulation produces a hippocampal formation theta rhythm in anesthetized rats. Brain Res. 1675, 41–50 (2017).

148. Broncel, A., Bocian, R., Kłos-Wojtczak, P. & Konopacki, J. Some technical issues of vagal nerve stimulation. An approach using a hippocampal formation theta rhythm. Brain Res. Bull. 140, 402–410 (2018).

149. Broncel, A., Bocian, R., Kłos-Wojtczak, P. & Konopacki, J. Medial septal cholinergic mediation of hippocampal theta rhythm induced by vagal nerve stimulation. PLOS ONE 13, e0206532 (2018).

150. Buckner, R. L. The serendipitous discovery of the brain’s default network. NeuroImage 62, 1137–1145 (2012).

151. Binder, J. R., Desai, R. H., Graves, W. W. & Conant, L. L. Where Is the Semantic System? A Critical Review and Meta-Analysis of 120 Functional Neuroimaging Studies. Cereb. Cortex 19, 2767–2796 (2009).

152. Levinthal, D. J. & Strick, P. L. The Motor Cortex Communicates with the Kidney. J. Neurosci. 32, 6726–6731 (2012).

153. Levinthal, D. J. & Strick, P. L. Multiple areas of the cerebral cortex influence the stomach. Proc. Natl. Acad. Sci. 117, 13078–13083 (2020).

154. Barnett, A. J. et al. Intrinsic connectivity reveals functionally distinct cortico-hippocampal networks in the human brain. PLOS Biol. 19, e3001275 (2021).

155. Cenquizca, L. A. & Swanson, L. W. Spatial organization of direct hippocampal field CA1 axonal projections to the rest of the cerebral cortex. Brain Res. Rev. 56, 1–26 (2007).

156. Chrobak, J. J. & Amaral, D. G. Entorhinal cortex of the monkey: VII. Intrinsic connections. J. Comp. Neurol. 500, 612–633 (2007).

157. Canto, C. B., Wouterlood, F. G. & Witter, M. P. What Does the Anatomical Organization of the Entorhinal Cortex Tell Us? Neural Plast. 2008, e381243 (2008).

158. Kerr, K. M., Agster, K. L., Furtak, S. C. & Burwell, R. D. Functional neuroanatomy of the parahippocampal region: The lateral and medial entorhinal areas. Hippocampus 17, 697–708 (2007).

159. Navarro Schröder, T., Haak, K. V., Zaragoza Jimenez, N. I., Beckmann, C. F. & Doeller, C. F. Functional topography of the human entorhinal cortex. eLife 4, e06738 (2015).

160. Blessing, E. M., Beissner, F., Schumann, A., Brünner, F. & Bär, K.-J. A data-driven approach to mapping cortical and subcortical intrinsic functional connectivity along the longitudinal hippocampal axis. Hum. Brain Mapp. 37, 462–476 (2016).

161. Boccia, M., Sulpizio, V., Nemmi, F., Guariglia, C. & Galati, G. Direct and indirect parietomedial temporal pathways for spatial navigation in humans: evidence from resting-state functional connectivity. Brain Struct. Funct. 222, 1945–1957 (2017).

162. Plachti, A. et al. Multimodal Parcellations and Extensive Behavioral Profiling Tackling the Hippocampus Gradient. Cereb. Cortex 29, 4595–4612 (2019).

163. Robinson, J. L. et al. Neurofunctional topography of the human hippocampus. Hum. Brain Mapp. 36, 5018–5037 (2015).

164. Adnan, A. et al. Distinct hippocampal functional networks revealed by tractography-based parcellation. Brain Struct. Funct. 221, 2999–3012 (2016).

165. Dickerson, B. C. & Eichenbaum, H. The Episodic Memory System: Neurocircuitry and Disorders. Neuropsychopharmacology 35, 86–104 (2010).

166. Libby, L. A., Ekstrom, A. D., Ragland, J. D. & Ranganath, C. Differential Connectivity of Perirhinal and Parahippocampal Cortices within Human Hippocampal Subregions Revealed by High-Resolution Functional Imaging. J. Neurosci. 32, 6550–6560 (2012).

167. Poppenk, J., Evensmoen, H. R., Moscovitch, M. & Nadel, L. Long-axis specialization of the human hippocampus. Trends Cogn. Sci. 17, 230–240 (2013).

168. Ranganath, C. & Ritchey, M. Two cortical systems for memory-guided behaviour. Nat. Rev. Neurosci. 13, 713–726 (2012).

169. Ritchey, M. & Cooper, R. A. Deconstructing the Posterior Medial Episodic Network. Trends Cogn. Sci. (2020) doi:10.1016/j.tics.2020.03.006.

170. Robin, J. & Moscovitch, M. Details, gist and schema: hippocampal–neocortical interactions underlying recent and remote episodic and spatial memory. Curr. Opin. Behav. Sci. 17, 114–123 (2017).

171. Poppenk, J. & Moscovitch, M. A Hippocampal Marker of Recollection Memory Ability among Healthy Young Adults: Contributions of Posterior and Anterior Segments. Neuron 72, 931–937 (2011).

172. Grady, C. L. Meta-analytic and functional connectivity evidence from functional magnetic resonance imaging for an anterior to posterior gradient of function along the hippocampal axis. Hippocampus 30, 456–471 (2020).

173. Paleja, M., Girard, T. A., Herdman, K. A. & Christensen, B. K. Two distinct neural networks functionally connected to the human hippocampus during pattern separation tasks. Brain Cogn. 92, 101–111 (2014).

174. Rochefort, C., Lefort, J. & Rondi-Reig, L. The cerebellum: a new key structure in the navigation system. Front. Neural Circuits 7, (2013).

175. Bohne, P., Schwarz, M. K., Herlitze, S. & Mark, M. D. A New Projection From the Deep Cerebellar Nuclei to the Hippocampus via the Ventrolateral and Laterodorsal Thalamus in Mice. Front. Neural Circuits 13, (2019).

176. Arrigo, A. et al. Constrained spherical deconvolution analysis of the limbic network in human, with emphasis on a direct cerebello-limbic pathway. Front. Hum. Neurosci. 8, (2014).

177. Baldassano, C. et al. Discovering Event Structure in Continuous Narrative Perception and Memory. Neuron 95, 709–721.e5 (2017).

178. Van Overwalle, F., Manto, M., Leggio, M. & Delgado-García, J. M. The sequencing process generated by the cerebellum crucially contributes to social interactions. Med. Hypotheses 128, 33–42 (2019).

179. Zacks, J. M. et al. Event perception: A mind-brain perspective. Psychol. Bull. 133, 273–293 (2007).

180. Hasselmo, M. E. What is the function of hippocampal theta rhythm?—Linking behavioral data to phasic properties of field potential and unit recording data. Hippocampus 15, 936–949 (2005).

181. Barrett, L. F., Quigley, K. S. & Hamilton, P. An active inference theory of allostasis and interoception in depression. Philos. Trans. R. Soc. B Biol. Sci. 371, 20160011 (2016).

182. Smith, R., Badcock, P. & Friston, K. J. Recent advances in the application of predictive coding and active inference models within clinical neuroscience. Psychiatry Clin. Neurosci. 75, 3–13 (2021).

183. Glasser, M. F. et al. The minimal preprocessing pipelines for the Human Connectome Project. NeuroImage 80, 105–124 (2013).

184. Smith, S. M. et al. Resting-state fMRI in the Human Connectome Project. NeuroImage 80, 144–168 (2013).

185. Li, J. et al. Global signal regression strengthens association between resting-state functional connectivity and behavior. NeuroImage 196, 126–141 (2019).

