## Supplementary Methods for "Functional connectivity gradients as a common neural architecture for predictive processing in the human brain"

**Datasets.** *Human Connectome Project (HCP)*. We analyzed the fMRI data collected at wakeful rest from 1,003 participants as part of the HCP WU-Minn Consortium [1] ($M_{age}$ = 28.71, $SD_{age}$ = 3.71, 470 males, 533 females; four 15 min runs per participant) included in the HCP1200 2017 data release. In particular, we took advantage of the group average preprocessed dense functional connectome data identifying correlation values between all cortical vertices and subcortical voxels. A full description of the data preprocessing pipelines implemented by the HCP is discussed elsewhere [2,3]. Briefly, each participant's fMRI data underwent gradient distortion correction, EPI distortion correction, motion correction, spatial co-registration to structural reference, spatial normalization to template volumetric space, resampling to template surface space, volumetric and surface smoothing with a 2 mm Gaussian kernel, and were submitted to independent component analysis (ICA) for further artifact removal [4,5]. For inter-subject registration, feature-based alignment and the Multimodal Surface Matching Algorithm (MSMAll) were implemented [6,7]. Each dataset was temporally demeaned and had variance normalization applied [8] and submitted to the group-level principal component analysis (PCA). The output of the group-PCA (the top 4,500 weighted spatial eigenvectors) are then renormalized, eigenvalue-reweighted, and correlated to form the group average dense connectome data (91,282 × 91,282 entries). We did not perform any further preprocessing on these data beyond what had already been implemented by the HCP.

*Brain Genomics Superstruct Project (GSP).* To validate the findings from the HCP dataset, we additionally analyzed structural and functional MRI data part of the Brain Genomics Superstruct Project (GSP). A comprehensive description of the GSP dataset is discussed elsewhere [9,10]. Briefly, this dataset includes 1,139 participants ($M_{age}$ = 21.24, $SD_{age}$ = 2.70, 467 males, 672 females) who had undergone one structural scan (T1-weighted multi-echo MPRAGE, 1.2 mm isotropic voxels) and two 6 min functional runs at wakeful rest (gradient-echo EPI sequence, 3 mm isotropic voxels) using 3T Siemens Tim Trio Scanners.

**MRI data preprocessing.** Each participant's structural data underwent intensity normalization, skull stripping, and an automated segmentation of cerebral white matter to locate the gray/white boundary via the FreeSurfer image analysis suite (v6.0), which is documented and freely available for download online (http://surfer.nmr.mgh.harvard.edu/). Defects in the surface topology were corrected [11], and the gray/white boundary was deformed outward using an algorithm designed to obtain an explicit representation of the pial surface. Each participant's cortical surface mesh was registered to a common spherical coordinate system [12,13]. Preprocessing of functional data was performed using the surface-based pipeline developed by Yeo and colleagues [14,15] using a combination of FreeSurfer [16], FSL [17], and Advanced Normalization Tools (ANTs) [18] routines as well as additional MATLAB functions. This pipeline consisted of the following preprocessing steps: Removal of the first four frames, slice timing correction, motion correction with rigid body translation and rotation, motion outlier detection, functional-to-structural co-registration via boundary-based registration [19], nuisance regression, interpolation of censored frames with Lomb-Scargle periodogram [20], and band-pass filtering [0.009, 0.08 Hz]. Volumetric data were then projected onto the FreeSurfer fsaverage6 surface space (~2 mm vertex spacing) followed by surface-constrained smoothing with a 2 mm Gaussian kernel. Subcortical voxels were resampled to the MNI152 template space (2 mm isotropic resolution) and volumetrically smoothed with a 2 mm Gaussian kernel.

We estimated framewise displacement (FD) [21] and root-mean-square of voxel-wise differentiated signal (DVARS) [22] using fsl_motion_outliers [23]. Volumes with FD > 0.2 mm or DVARS > 50 were marked as outliers (censored frames), following the criteria used by previous studies [14,15]. One frame before and two frames after these outlier volumes, as well as uncensored segments of BOLD data lasting fewer than five contiguous volumes, were also flagged as censored frames [24]. BOLD runs with more than 50% of the volumes labeled as censored frames were discarded.

To account for the effect of confounding variables, we performed linear regression separately for each BOLD run with multiple nuisance regressors, including (1) a vector of ones and linear trend, (2) six motion parameters, (3) averaged white matter signal, (4) averaged ventricular signal, along with the first-order temporal derivatives of (2), (3), and (4). The white matter mask for each participant was derived from FreeSurfer's segmentation of their structural image, followed by three rounds of erosion before resampling to their native BOLD space. The ventricular mask was obtained similarly, but only with one round of erosion. In the event that there were fewer than 100 voxels after a round of erosion, no further erosion was performed. Regression coefficients were computed without censored frames [20]. To maintain consistency with the HCP dataset, we resampled the denoised BOLD time series data in the fsaverage6 space to fs_LR 32k space, after which these surface data were combined with the volumetric data to form a single whole-brain time series file per run.

From the original pool of 1,139 participants with two BOLD runs, we discarded 12 participants who had at least one run with more than 50% of the volumes labeled as censored frames. We additionally discarded 25 participants for whom surface resampling resulted in fewer vertices/voxels in at least one of the runs than the rest of the participants. The final GSP dataset analyzed in the current study thus consisted of 1,102 individuals. For each participant, we concatenated the two BOLD runs and computed the Pearson's correlation coefficient between every pair of vertices/voxels, which was standardized via Fisher's r-to-z transformation. Individual-level Z maps were averaged across all 1,102 participants to yield the group average dense connectome data.

**Diffusion map embedding.** We derived functional connectivity gradients of the cerebral cortex, the cerebellum, and the hippocampus using diffusion map embedding [25,26]. Diffusion map embedding is a non-linear data dimensionality reduction technique that enables an analysis of similarity structure in functional connectivity patterns in a large number of data points (e.g., vertices/voxels) by identifying a set of low-dimensional manifolds (i.e., gradients) capturing principal dimensions of spatial variation in connectivity. Based on the group average dense connectome data, we first derived functional connectivity matrices between (1) all cerebral cortical vertices (cortico-cortical symmetric matrix), (2) all cerebellar voxels and all cerebral cortical vertices (cerebello-cortical asymmetric matrix), and (3) all unilateral hippocampal voxels and all cerebral cortical vertices (hippocampo-cortical asymmetric matrix). We defined the voxels belonging to the cerebellum and the hippocampus based on a probabilistic cerebellar atlas [27] and the Harvard-Oxford subcortical structural atlas [28,29], respectively (both thresholded at 50%).

We converted each functional connectivity matrix back to Pearson's r values using a hyperbolic tangent function and applied row-wise thresholding to retain the top 10% connections, with all other connections set to zero. To characterize the relationship (i.e., similarity) in functional connectivity between a given pair of cerebral cortical vertices or cerebellar/hippocampal voxels, we computed a non-negative square symmetric affinity matrix for each functional connectivity matrix. In keeping with the previous investigations [30,31], we opted to use cosine similarity to characterize the similarity structure in functional connectivity for the cortico-cortical and cerebello-cortical matrices and normalized angle similarity for the hippocampo-cortical matrices [32,33]. Finally, we used these affinity matrices as input to diffusion map embedding, which yielded ten gradients per affinity matrix identifying the dominant dimensions of spatial variation in the cerebral cortex, the cerebellum, and the left and right hippocampus in terms of their functional connectivity with (the rest of) the cerebral cortex. Given sign indeterminacy inherent in the use of this algorithm, we visually inspected each gradient and inverted the sign of vertex-wise or voxel-wise gradient values where appropriate (e.g., for consistency between the left and right hippocampus).

**Interpretation of functional connectivity gradients.** We performed post hoc characterization of the functional gradients identified via diffusion map embedding at various levels to interpret their significance. For each of the cerebral cortical gradients and gradient-weighted functional connectivity maps of the cerebellum and the hippocampus (see below), we characterized the distribution of gradient values across the seven canonical functional networks of the cerebral cortex [10]. For the hippocampal gradients, we characterized and compared the distribution of gradient values between major hippocampal subfields. To do this, we first performed automatic segmentation of hippocampal subfields on a T1-weighted structural image in the MNI152 space

[34,35]. This procedure generated binary ROIs of the subiculum, CA1-3, and CA4-DG [36], which were subsequently down-sampled to the resolution of functional data.

To interpret the cerebellar and hippocampal gradients in terms of their relations to the cerebral cortex, we calculated functional connectivity maps between these structures and the cerebral cortex weighted as a factor of voxel-wise gradient values, following previous studies of functional connectivity gradients [37,38]. We performed this procedure separately for the left and right hippocampus, then the resulting weighted connectivity maps corresponding to the same gradient were averaged across hemispheres. This decision was justified by the high correlation of weighted functional connectivity maps observed across hemispheres ($G_1$: r = .93, $G_2$: r = .94). To statistically assess the correspondence between the cerebral cortical gradients and the gradient-weighted functional connectivity maps of the cerebellum and the hippocampus, we computed vertex-wise Spearman's rank correlations. We conducted non-parametric spin tests to derive statistical significance of each association while controlling for autocorrelations [39].

**Gradient-informed triangulation of intrinsic functional connectivity between the cerebral cortex, the cerebellum, and the hippocampus.** To further demonstrate the correspondence between the cerebral cortical, cerebellar, and hippocampal gradients in terms of connectivity, we performed a seed-based functional connectivity analysis. This analysis was performed separately for two sets of the cerebral cortical, cerebellar, and hippocampal connectivity gradients identified as being maximally correlated with one another and corresponding to unique predictive processing gradients (see **Results** and **Fig. 3c**). Our analysis of seed-based functional connectivity proceeded as follows: (1) For each gradient derived from diffusion map embedding, we identified the vertices/voxels with the top and bottom 10% gradient values, which were defined as seed ROIs (**Fig. 4A-L**). (2) For each seed ROI in a given structure, we computed the mean BOLD activity time course based on all vertices or voxels and correlated it with the time course of every vertex or voxel in the other two structures. This step yielded two functional connectivity maps per seed ROI (e.g., each cerebral cortical ROI had one connectivity map representing its functional connectivity with all voxels in the cerebellum and another connectivity map representing its functional connectivity with all voxels in the hippocampus). In other words, this step resulted in four functional connectivity maps per structure (e.g., the cerebral cortex had functional connectivity maps seeded in the top 10% voxels of the cerebellum, the top 10% voxels of the hippocampus, the bottom 10% voxels of the cerebellum, and the bottom 10% voxels of the hippocampus, all of which corresponded to the same gradient). (3) For each structure, we identified the vertices/voxels that were reliably connected to the subregions of the other two structures that anchored the corresponding end of their respective gradients. This step involved calculation of functional connectivity maps in a given structure through a combination of binarization and inclusive masking of the contributing maps as well as proportional thresholding. For instance, to identify the cerebral cortical vertices functionally connected to both the cerebellar and hippocampal voxels representing the top 10% of their respective gradients, we extracted the mean BOLD activity time course of the top 10% voxels in the cerebellum and the hippocampus and correlated them separately with that of every cerebral cortical vertex. We then binarized these connectivity maps at $Z > 0$ and calculated their intersection, thus retaining only the cerebral cortical vertices showing positive connectivity with both of the other two structures. Each cerebral cortical vertex within this restricted set was assigned the weighted average functional connectivity value to account for the difference in the number of voxels between the cerebellar and hippocampal seed ROIs. Finally, we thresholded the resulting connectivity maps at 50th percentile. This procedure was repeated for each structure and for each end (top vs. bottom 10%) of the gradients. Altogether, this analysis allowed us to examine the extent to which the subregions of the cerebral cortex, the cerebellum, and the hippocampus anchoring the corresponding end of their respective gradients showed preferential functional connectivity with each other.

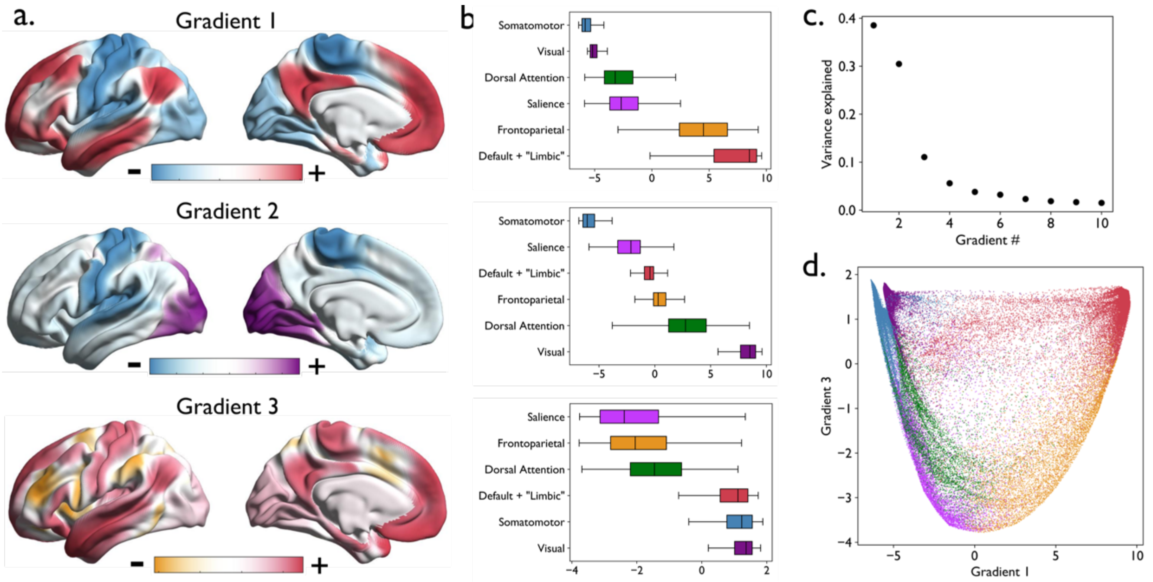


**Fig. S1.** The principal gradients of human cerebral cortex based on the GSP data ($n$ = 1,102). (a) The three most dominant gradients projected onto an inflated cortical surface (left hemisphere only). These gradients replicate previous findings and identify model-error (Gradient 1), visual-sensorimotor (Gradient 2), and model-precision (Gradient 3) gradients. (b) Box plots show the median and distribution of gradient values separately for each of the canonical functional network [10]. The networks are ordered by the mean value. Vertices belonging with the so-called default mode and "limbic" networks are shown in the same color, as these networks are not always distinguished in the literature [14] and both contain agranular, limbic tissue [40]. (c) A scree plot showing the proportion of variance explained by each of the ten gradients derived from diffusion map embedding. (d) A scatterplot depicting the relationship between Gradient 1 and Gradient 3, where each dot represents a cerebral cortical vertex color-coded by the corresponding network assignment.

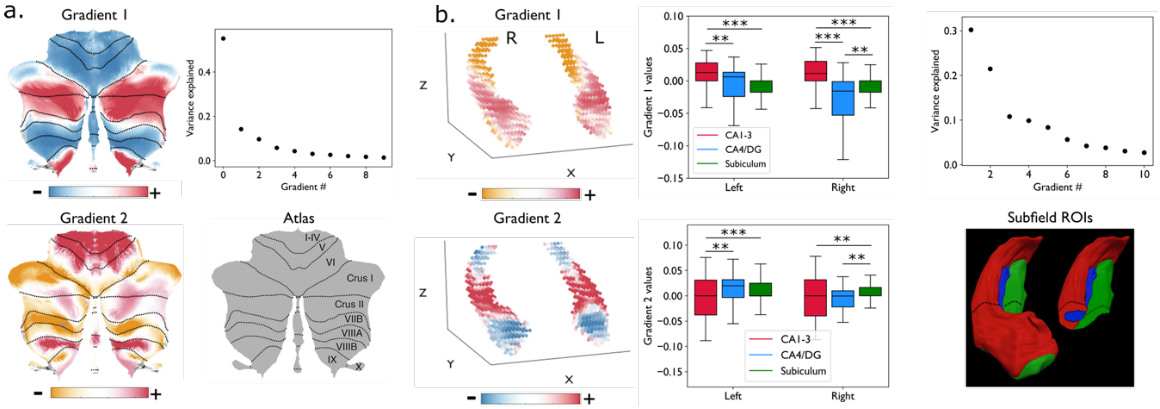


**Fig. S2.** The principal gradients of human cerebellum and hippocampus based on the GSP data (*n* = 1,102). (a) The two most dominant gradients of the cerebellum replicated previous findings [30]. (b) The most dominant gradient of the hippocampus replicated previous findings [33] and identified an anterior-posterior dissociation along the longitudinal axis, which also differentiated the major hippocampal subfields. The second most dominant gradient was also consistent with differences by hippocampal microstructure, with the subiculum exhibiting highest gradient values overall compared with the other two subregions, with the exception of the left CA4-DG where the difference did not reach significance at a conventional threshold of $\alpha = .05$ ($p < .12$). Box plots show the median and distribution of $G_2$ values per subfields separately for each hemisphere. Asterisks denote significant (***$p < .001$, **$p \leq .01$) differences relative to the other two subfields. DG = dentate gyrus. A figure illustrating the hippocampal subfields within the right hippocampus (red = CA1-3, blue = CA4-DG, green = subiculum) was reproduced from [36] with permission.

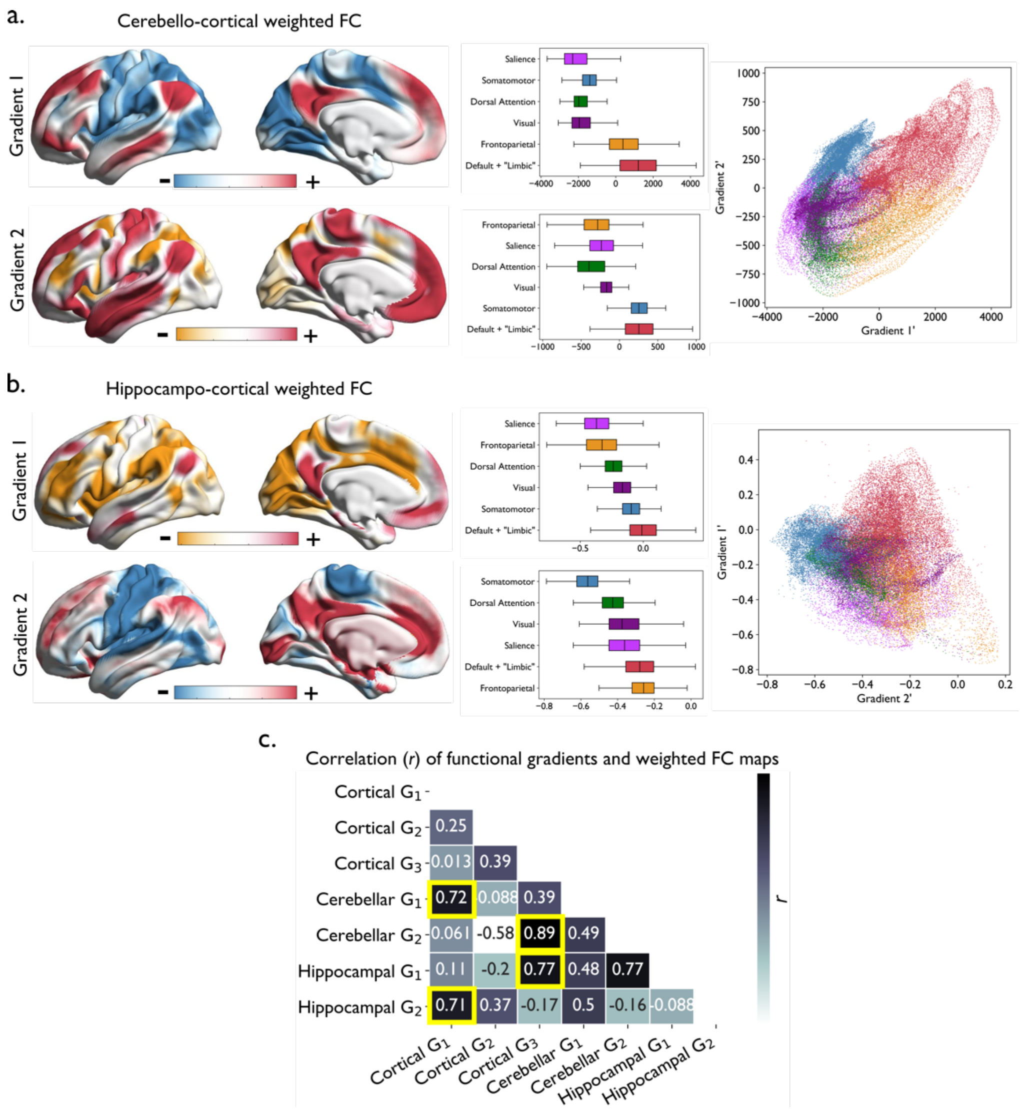


**Fig. S3.** Gradient-weighted functional connectivity maps of the cerebellum and the hippocampus. (a) Cerebellar $G_1$ captured a dissociation in functional connectivity most consistent with cortical $G_1$, from ensembles including the default mode and frontoparietal networks to those including the exteroceptive sensory and salience networks (i.e., the model-error gradient). In contrast, cerebellar $G_2$ captured a dissociation in connectivity most consistent with cortical $G_3$, from ensembles including the default mode and exteroceptive networks to those including the frontoparietal and salience networks (i.e., the model-precision gradient). Box plots show the median and distribution of gradient-weighted functional connectivity values separately for each of the canonical functional network [10]. The networks are ordered by the mean value. Vertices belonging with the default and "limbic" networks are shown in the same color, as these networks are not always distinguished in the literature [14]. A scatterplot depicts the relationship between the gradient-weighted functional connectivity maps (Gradient 1' and Gradient 2'), where each dot represents a cortical vertex color-coded by the corresponding network assignment. (b) Hippocampal $G_1$ captured a dissociation in functional connectivity most consistent with cortical $G_3$, whereas hippocampal $G_2$ captured a dissociation in connectivity most consistent with cortical $G_1$. (c) A similarity matrix illustrating the magnitude (Spearman's r) of correlation between cortical gradients and gradient-weighted functional connectivity maps of the cerebellum and the hippocampus. Highlighted in yellow are the strongest cortico-cerebellar or cortico-hippocampal associations identifying the correspondence of gradients between a pair of structures. *p*-values associated with the entire correlation matrix are shown in **Fig. S5**. FC, functional connectivity.

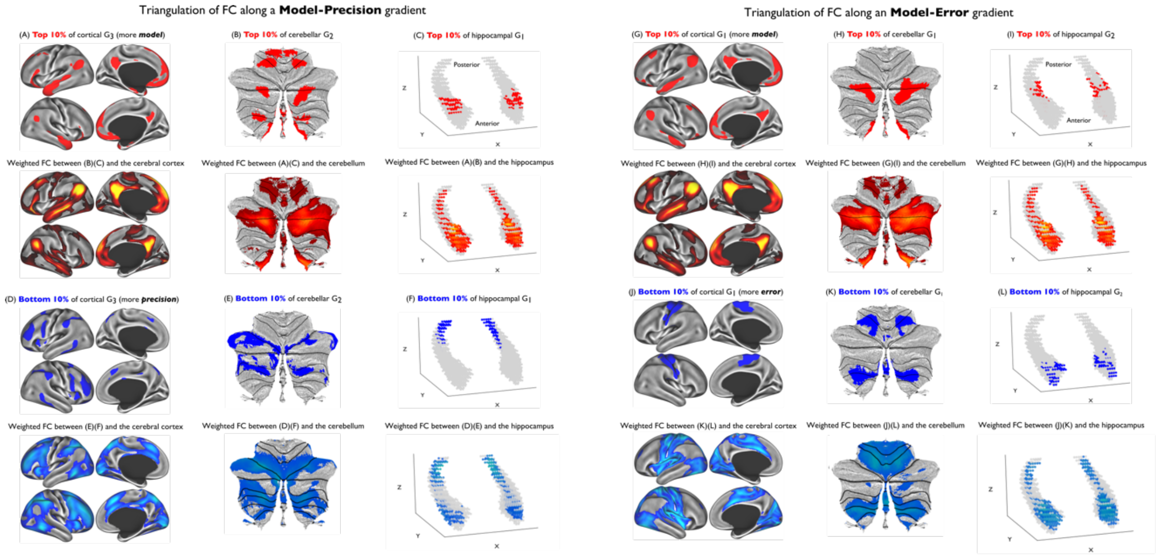


**Fig. S4.** Triangulation of intrinsic functional connectivity between the cerebral cortex, the cerebellum, and the hippocampus along the cortical representation-modulation (left) and the cortical model-periphery (right) gradients in the GSP ($n$ = 1,102) data. These analyses were performed using the seed ROIs derived from the HCP dataset (A-F). Functional connectivity maps shown here for a given structure were calculated using the GSP data through a combination of binarization and inclusive masking of the contributing maps as well as proportional thresholding (see **Methods**). We expect, and indeed observe, that there is remarkable spatial overlap between a given seed ROI (e.g., the top 10% of the vertices in cortical representation-modulation $G_3$) and areas of the same structure functionally connected to parts of the other two structures anchoring the same end of the gradient (e.g., the top 10% of the voxels in cerebellar $G_2$ and hippocampal $G_1$).

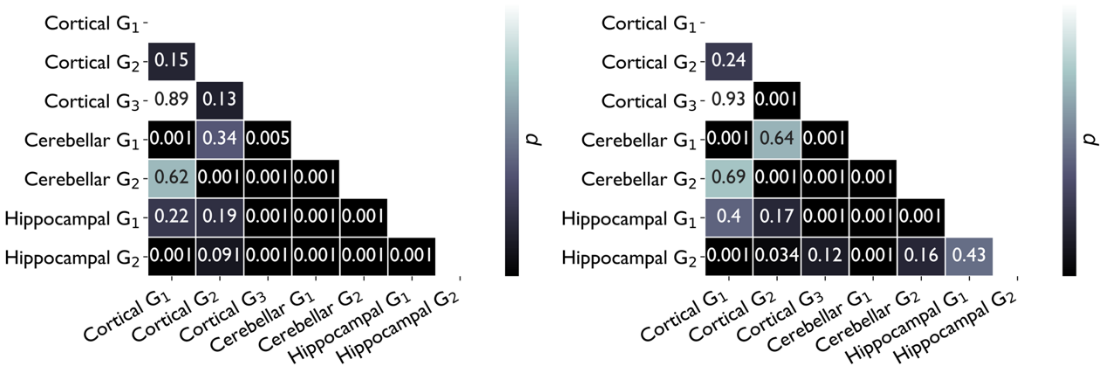


**Fig. S5.** Statistical significance ($p$-values) associated with vertex-wise correlation of cortical gradients and cortical functional connectivity maps weighted by a factor of cerebellar and hippocampal gradients. Left, HCP dataset; right, GSP dataset.

**References**


1. Van Essen, D. C. *et al.* The WU-Minn Human Connectome Project: An overview. *NeuroImage* **80**, 62–79 (2013).
2. Glasser, M. F. *et al.* The minimal preprocessing pipelines for the Human Connectome Project. *NeuroImage* **80**, 105–124 (2013).
3. Smith, S. M. *et al.* Resting-state fMRI in the Human Connectome Project. *NeuroImage* **80**, 144–168 (2013).
4. Griffanti, L. *et al.* ICA-based artefact removal and accelerated fMRI acquisition for improved resting state network imaging. *NeuroImage* **95**, 232–247 (2014).
5. Salimi-Khorshidi, G. *et al.* Automatic denoising of functional MRI data: Combining independent component analysis and hierarchical fusion of classifiers. *NeuroImage* **90**, 449–468 (2014).
6. Glasser, M. F. *et al.* A multi-modal parcellation of human cerebral cortex. *Nature* **536**, 171–178 (2016).
7. Robinson, E. C. *et al.* MSM: A new flexible framework for Multimodal Surface Matching. *NeuroImage* **100**, 414–426 (2014).
8. Beckmann, C. F. & Smith, S. M. Probabilistic independent component analysis for functional magnetic resonance imaging. *IEEE Trans. Med. Imaging* **23**, 137–152 (2004).
9. Holmes, A. J. *et al.* Brain Genomics Superstruct Project initial data release with structural, functional, and behavioral measures. *Sci. Data* **2**, 1–16 (2015).
10. Yeo, B. T. T. *et al.* The organization of the human cerebral cortex estimated by intrinsic functional connectivity. *J. Neurophysiol.* **106**, 1125–1165 (2011).
11. Fischl, B., Liu, A. & Dale, A. M. Automated manifold surgery: constructing geometrically accurate and topologically correct models of the human cerebral cortex. *IEEE Trans. Med. Imaging* **20**, 70–80 (2001).
12. Fischl, B., Sereno, M. I., Tootell, R. B. H. & Dale, A. M. High-resolution intersubject averaging and a coordinate system for the cortical surface. *Hum. Brain Mapp.* **8**, 272–284 (1999).
13. Fischl, B., Sereno, M. I. & Dale, A. M. Cortical Surface-Based Analysis: II: Inflation, Flattening, and a Surface-Based Coordinate System. *NeuroImage* **9**, 195–207 (1999).
14. Kong, R. *et al.* Spatial Topography of Individual-Specific Cortical Networks Predicts Human Cognition, Personality, and Emotion. *Cereb. Cortex* **29**, 2533–2551 (2019).
15. Li, J. *et al.* Global signal regression strengthens association between resting-state functional connectivity and behavior. *NeuroImage* **196**, 126–141 (2019).
16. Fischl, B. FreeSurfer. *NeuroImage* **62**, 774–781 (2012).
17. Jenkinson, M., Beckmann, C. F., Behrens, T. E. J., Woolrich, M. W. & Smith, S. M. FSL. *NeuroImage* **62**, 782–790 (2012).
18. Avants, B. B. *et al.* A reproducible evaluation of ANTs similarity metric performance in brain image registration. *NeuroImage* **54**, 2033–2044 (2011).
19. Greve, D. N. & Fischl, B. Accurate and robust brain image alignment using boundary-based registration. *NeuroImage* **48**, 63–72 (2009).
20. Power, J. D. *et al.* Methods to detect, characterize, and remove motion artifact in resting state fMRI. *NeuroImage* **84**, 320–341 (2014).
21. Jenkinson, M., Bannister, P., Brady, M. & Smith, S. Improved Optimization for the Robust and Accurate Linear Registration and Motion Correction of Brain Images. *NeuroImage* **17**, 825–841 (2002).
22. Power, J. D., Barnes, K. A., Snyder, A. Z., Schlaggar, B. L. & Petersen, S. E. Spurious but systematic correlations in functional connectivity MRI networks arise from subject motion. *NeuroImage* **59**, 2142–2154 (2012).
23. Smith, S. M. *et al.* Advances in functional and structural MR image analysis and implementation as FSL. *NeuroImage* **23**, S208–S219 (2004).
24. Gordon, E. M. *et al.* Generation and Evaluation of a Cortical Area Parcellation from Resting-State Correlations. *Cereb. Cortex* **26**, 288–303 (2016).
25. Coifman, R. R. & Lafon, S. Diffusion maps. *Appl. Comput. Harmon. Anal.* **21**, 5–30 (2006).
26. Vos de Wael, R. *et al.* BrainSpace: a toolbox for the analysis of macroscale gradients in neuroimaging and connectomics datasets. *Commun. Biol.* **3**, 1–10 (2020).
27. Diedrichsen, J., Balsters, J. H., Flavell, J., Cussans, E. & Ramnani, N. A probabilistic MR atlas of the human cerebellum. *NeuroImage* **46**, 39–46 (2009).

28. Desikan, R. S. *et al.* An automated labeling system for subdividing the human cerebral cortex on MRI scans into gyral based regions of interest. *NeuroImage* **31**, 968–980 (2006).
29. Frazier, J. A. *et al.* Structural Brain Magnetic Resonance Imaging of Limbic and Thalamic Volumes in Pediatric Bipolar Disorder. *Am. J. Psychiatry* **162**, 1256–1265 (2005).
30. Guell, X., Schmahmann, J. D., Gabrieli, J. D. & Ghosh, S. S. Functional gradients of the cerebellum. *eLife* **7**, e36652 (2018).
31. Margulies, D. S. *et al.* Situating the default-mode network along a principal gradient of macroscale cortical organization. *Proc. Natl. Acad. Sci.* **113**, 12574–12579 (2016).
32. Li, Q. *et al.* Topographic profiling of memory-related pattern separation processes in humans. *bioRxiv* 2020.06.22.165290 (2020) doi:10.1101/2020.06.22.165290.
33. Vos de Wael, R. *et al.* Anatomical and microstructural determinants of hippocampal subfield functional connectome embedding. *Proc. Natl. Acad. Sci.* **115**, 10154–10159 (2018).
34. Manjón, J. V. & Coupé, P. volBrain: An Online MRI Brain Volumetry System. *Front. Neuroinformatics* **10**, (2016).
35. Romero, J. E., Coupé, P. & Manjón, J. V. HIPS: A new hippocampus subfield segmentation method. *NeuroImage* **163**, 286–295 (2017).
36. Kulaga-Yoskovitz, J. *et al.* Multi-contrast submillimetric 3 Tesla hippocampal subfield segmentation protocol and dataset. *Sci. Data* **2**, 1–9 (2015).
37. Guell, X. *et al.* Functional Territories of Human Dentate Nucleus. *Cereb. Cortex* (2020) doi:10.1093/cercor/bhz247.
38. Zhang, J. *et al.* Intrinsic functional connectivity is organized as three interdependent gradients. *Sci. Rep.* **9**, 1–14 (2019).
39. Alexander-Bloch, A. F. *et al.* On testing for spatial correspondence between maps of human brain structure and function. *NeuroImage* **178**, 540–551 (2018).
40. Kleckner, I. R. *et al.* Evidence for a large-scale brain system supporting allostasis and interoception in humans. *Nat. Hum. Behav.* **1**, 1–14 (2017).
